# Protamine lacking piscine spermatozoa are transcriptionally active

**DOI:** 10.1101/2021.11.05.467500

**Authors:** Júlia Castro-Arnau, François Chauvigné, Jessica Gómez-Garrido, Anna Esteve-Codina, Marc Dabad, Tyler Alioto, Roderick Nigel Finn, Joan Cerdà

## Abstract

Transcriptional quiescence of post-meiotic spermatozoa associated with protamine-mediated chromatin condensation is widely recognized in animals. How sperm acquire the extratesticular maturational competence to move and fertilize the egg is therefore thought to occur via non-transcriptional mechanisms. Here, using transcriptional profiling during spermatozoon differentiation in a fish that does not condense chromatin with protamines, we uncover spatially distinct roles of the GnRH receptor and PDGF signaling pathways between the somatic epithelia of the extratesticular ducts and the maturing spermatozoa. In vitro induction and inhibition experiments demonstrate that the endocrine signaling pathways are conserved in different lineages of fish and activate de novo transcription of spermatozoon genes required for the acquisition of full motility. These experiments further confirmed that mitochondrial translation is important for sperm maturation in anamniotes as in amniotes, but that transcriptional quiescence of post-meiotic spermatozoa is not a pan vertebrate phenomenon. On the contrary, the data show that the identified signal transduction pathways between the soma and the sperm upregulate effector genes essential for maturational competence and male fertility.

## Introduction

Vertebrate spermatogenesis proceeds in a multistage process from mitotic expansion of spermatogonial stem cells to form primary spermatocytes (spermatocytogenesis) through two meiotic divisions to form spermatids (spermatidogenesis) and a tertiary phase of differentiation (spermiogenesis) to form the highly polarized spermatozoa that retain a recombined haploid genome (***de Kretser et al., 1998; Schulz et al., 2010; Nishimura and L’Hernault, 2017***). This is regardless of whether the germ cells develop in cysts in anamniotes (fishes and amphibians) or in the acystic epithelial lining of the seminiferous tubules in amniotes (reptiles, birds and mammals) (***Yoshida, 2016***). At the culmination of testicular spermatogenesis however, the fully differentiated spermatozoa are typically not capable of fertilizing the egg (***Nixon et al., 2020; Pérez, 2020***). They require a maturational phase, which confers the physiological ability to move, recognize and penetrate the egg (***Nixon et al., 2020; Pérez, 2020***). In most vertebrates, this process proceeds during sperm storage and transit through the extratesticular excurrent ducts (ETDs) and tubular systems that emanate from the testis (***Sullivan and Mieusset, 2016***). Such ETDs are thought to have evolved in the common ancestor of jawed vertebrates becoming ever more convoluted to form the epididymis in amniotes (***Jones, 2002***). Since humans are members of this latter group, considerable research has been invested to understand the epididymal regulation of sperm maturation and the aetiology of asthenozoospermia (***Sullivan and Mieusset, 2016***). By contrast, almost nothing is known of the molecular signaling pathways that regulate sperm maturation in anamniotes.

Both transcriptomic and proteomic studies in mammals have established that gene expression is highly segmented along the length of the epididymis (***Sullivan and Mieusset, 2016***; ***Belleannée et al., 2012***; ***Zhao et al., 2019***). Conversely, despite presenting hundreds of proteins and carrying thousands of RNAs of different types, the transcriptional and translational activities of the maturing spermatozoa are virtually silent (Fisher et al., 2012; ***Grunewald et al., 2005***; ***Ren et al., 2017***; ***Freitas et al., 2020***). Such quiescence is in stark contrast to the stellar transcriptional activity of the spermatogenic cells, which exceed all other cell types by expressing >80% of the protein coding genes in the genome (***Soumillon et al., 2013***; ***Xia et al., 2020***). The onset of transcriptional and translational quiescence occurs during the spermiogenic differentiation phase when large numbers of rRNAs are degraded, the cytoplasm and nucleoplasm are discarded, and the histones of the DNA-packing nucleosomes are gradually replaced by protamines (***Ren et al., 2017***; ***Rathke et al., 2014***). The many types of RNAs still present in the sperm are thus thought to be the remnants of the high transcriptional activity of the spermatocytic and spermatidogenic phases (***Ren et al., 2017***). Alternatively, many proteins and some types of RNAs may be delivered via epididymal exosomes-epididysomes (***James et al., 2020***), which partially solves the problem of the lack of cytoplasmic ribosomes for protein translation. Other studies suggest that mitochondrial ribosomal pathways, rather than the canonical cytoplasmic mechanisms, remain active and yield paternal factors that are important for sperm maturation, capacitation in the female reproductive tract, fertility and early zygotic development (***Gur and Breitbart, 2006***; ***Zhao et al., 2009***; ***Rajamanickam et al., 2017***; ***Zhu et al., 2019***). In all cases, however, *de novo* transcription in the maturing mammalian sperm is not considered to be a major source of RNAs or proteins.

Interestingly, not all vertebrate sperm retain protamines in their nuclei. Despite protamines first being discovered in fish, the Rhine salmon (***Miescher, 1874***), it has become evident that the spermatozoa of several lineages of anamniotes lack such highly arginine-enriched forms of the sperm nuclear basic proteins (SNBP) (***Ausió, 1999***; Eirin-López et al., 2006). Even when present, it has been shown that protamines may not be involved in spermatogenic nuclear condensation (***Shimizu et al., 2000***). It is thus not known whether transcriptional quiescence is a general feature of post-meiotic sperm maturation in vertebrates, and if not what role such late-stage transcription might play. To address these questions, we conducted transcriptome profiling of haploid germ cells and ejaculated spermatozoa from a species of fish, the gilthead seabream (*Sparus aurata*), which produces profuse amounts of sperm without protamines (***Kurtz et al., 2009***). Gene set enrichment analysis revealed the regulation of a high number of transcripts potentially involved in transcription, translation and chromatin organization in spermatozoa, as well as of several signaling pathways, of which the gonadotropin-releasing hormone receptor (GnRHR) and platelet-derived growth factor (PDGF) were amongst the most dominant. Experimental investigation of the origin of these signaling pathways uncovered their expression in sequential segments of the ETDs where their paracrine signaling induces the *de novo* transcription of genes in the post-meiotic spermatozoa. The importance of these mechanisms for sperm maturation in seabream, as well as in zebrafish (*Danio rerio*), as a model from a more ancestral teleost lineage that produces small volumes of sperm lacking protamines (***Wu et al., 2011***), was confirmed through motility tests in the presence of transcription and translation inhibitors. The present data thus uncover soma to germ cell signaling pathways during sperm maturation in vertebrates and reveal that post-meiotic spermatozoa may not remain transcriptionally silent. On the contrary, such late-stage transcriptional activation induced by ETD epithelial endocrine signaling upregulates gametic cell effector genes that are required for the acquisition of full sperm motility.

## Results

### Transcriptome profiling of haploid germ cells and mature spermatozoa

The changes in gene expression during the differentiation and maturation of seabream spermatozoa were investigated by whole-transcriptome RNA-seq of haploid germ cells (HGCs) and ejaculated (mature) spermatozoa (SPZ_EJ_). The HGCs were isolated by fluorescence-activated cell sorting (FACS), whereas SPZ_EJ_ were collected by manual stripping of naturally spermiating males. Flow cytometry of the extract from the seabream whole mature testis showed that the percentage of diploid and haploid cells reached 16% and 84% of the total cells, respectively (***Figure 1A***). The percentage of diploid cells was lower than expected because the centrifugation steps of the extract before cell sorting partially depleted this population. Flow cytometry identified two subpopulations of haploid cells based on their relative size and SYBR Green I fluorescence intensity: a subpopulation formed by spermatocytes II and spermatids (SPC II and SPD, respectively), which we refer here as HGCs, and another subpopulation formed by intratesticular spermatozoa (SPZ_I_) (***Figure 1A and B***). The percentage of HGCs and SPZ_I_ in the testicular extracts was of 34 ± 4% and 66 ± 4% (*n* = 15), respectively (***Figure 1B***).

**Figure 1.**
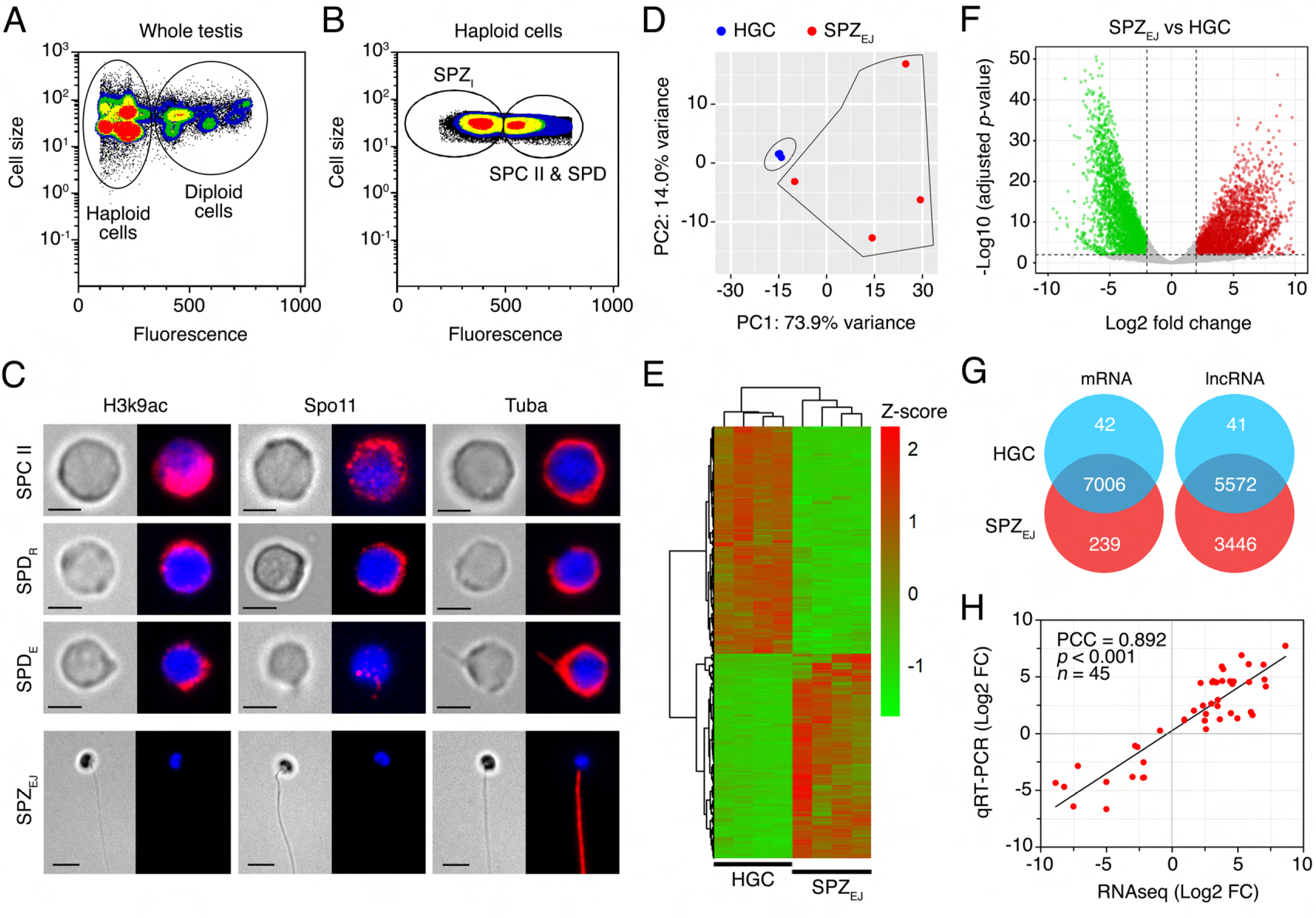
Transcriptome profiling of seabream haploid germ cells and ejaculated spermatozoa. (**A** and **B**) Representative flow cytometry plots of the seabream cell populations in the whole testis. In A, the populations of testicular haploid and diploid cells are encircled. In B, the different subpopulations of haploid germinal cells (HGC), corresponding to a mix of type II spermatocytes (SPC II) and spermatids (SPD), and intratesticular spermatozoa (SPZ_I_) are shown. (**C**) Representative immunostaining of Lys^9^ acetylated histone 3 (H3K9ac), meiotic recombination protein Spo11 and α-tubulin (Tuba) in sorted HGC and ejaculated spermatozoa (SPZ_EJ_). For each cell type the brightfield (left panels) and epifluorescence (right panels) images are shown. SPD_R_, round spermatids; SPD_E_, elongating spermatids. Scale bars, 2 and 5 µm. (**D**) Principal component analysis (PCA) using the top 500 most variable genes between HGC and SPZ_EJ_ (*n* = 4 pools) and the ‘rlog’ transformation of the counts. (**E**) Heatmap generated by unsupervised hierarchical clustering of RNAseq expression z-scores computed for the 7,287 differentially expressed genes (DEGs) (*p*-adj < 0.01; Log2 fold change > 1) between HGC and SPZ_EJ_. (**F**) Volcano plot representation of DEGs in the SPZ_EJ_ versus HGC comparison. The x-axis shows Log2 fold changes in expression and the y-axis the negative logarithm of their *p*-value to base 10. Red and green points mark the genes with significantly increased or decreased expression respectively in SPZ_EJ_ compared to HGC (FDR < 0.01). (**G**) Venn diagrams showing the number of common mRNAs and lncRNAs (in intersect region) which are differentially expressed between HGCs and SPZ_EJ_. (**H**) Validation of the RNAseq data by qRT-PCR. The plot represents the Pearson’s correlation analysis of DEGs in HGC and SPZ_EJ_ determined by RNAseq and qRT-PCR. The Pearson’s correlation coefficient (PCC) of the Log2 fold change analyzed by RNAseq (x-axis) and using qRT-PCR (y-axis), the *p*-value, and the number of DEGs analyzed are indicated. **Figure 1-source data 1** Data for PCA shown in D. **Figure 1-source data 2** Data for the heat map shown in E. **Figure 1-source data 3** Data for Volcano plot shown in F. **Figure 1-source data 4** Data on the validation of the RNAseq data by qRT-PCR.

Microscopic examination of the HGC-enriched population after FACS confirmed the presence of SPC II, and round and elongating SPD in this fraction (***Figure 1C***). Whole-mount immunostaining revealed strong expression of Lys^9^ acetylated histone 3 (H3K9ac) and meiotic recombination protein Spo11 in SPC II, which progressively decreased in round and elongating SPD, and completely vanished in SPZ_EJ_ (***Figure 1C***). Immunolocalization of α-tubulin (Tuba) showed that the protein was spread in the cytoplasm in SPC II and round SPD, became also detectable in the nascent flagellar region of elongating SPD, and was finally distributed along the flagellum of differentiated SPZ_EJ_ (***Figure 1C***). These observations indicate a high occurrence of meiotic recombination in SPC II and a progressive DNA condensation during the differentiation of SPC II into SPD and spermatozoa, which are conserved features during vertebrate germ cell development (***Kurtz et al., 2009***; ***Hazzouri et al., 2000***). Therefore, these data confirmed that the sorted population of cells from the mature seabream testis correspond to HGCs before differentiation into spermatozoa.

Four unstranded RNA libraries (replicates) for low-input RNA were subsequently constructed for each of the two HGC and SPZ_EJ_ cell types; each replicate being a pool of cells collected from three different males. Library sequencing rended 30-62 million reads per library comprising a yield of 5-10 Gb. From these data, we produced a new integrative *S. aurata* genome annotation before the RNA-seq analysis. This new annotation was carried out by re-annotating the available *S. aurata* reference genome (***Pauletto et al., 2018***), and by adding 202 *de novo* assembled transcripts that were not present in the genome assembly. In total, 31,501 protein-coding genes were annotated, which produced 57,396 transcripts (1.82 transcripts per gene) and encoded for 51,365 unique protein products. Functional labels were assigned to 62% of the annotated proteins. In addition, 165,898 non-coding transcripts were annotated, of which 159,925 are long non-coding RNA (lncRNA) genes and 5,973 correspond to short non-coding RNAs.

Principal component analysis (PCA) of the expression data showed that FACS-purified HGC and SPZ_EJ_ formed two relatively well-separated clusters, suggesting that the developmental stage has a large effect on the pattern of gene expression (***Figure 1D***). However, while the four HGC biological replicates clustered very close, those of SPZ_EJ_ were more distant, indicating a higher variability in the transcriptome of the SPZ_EJ_ replicates. Nevertheless, the RNA-seq analysis revealed a total of 7,287 differentially expressed genes (DEGs) (adjusted *p*-value < 0.01) between both cell types, of which nearly half (3,447) were upregulated in SPZ_EJ_ when compared to HGCs (***Figure 1E-G***). In addition, 239 transcripts were detected only in SPZ_EJ_ (***Figure 1G***). Finally, as previously reported in the human spermatozoon (Corral-Vázquez et al., 2021), we also found a high number of differentially expressed lncRNAs (9,059 sequences) of which 5,114 were upregulated and 3,446 unique in SPZ_EJ_ (***Figure 1G***).

The quality of the RNA-seq data and the reliability of the DEGs identified were validated on randomly selected 45 DEGs by real-time quantitative reverse transcription PCR (qRT-PCR) in three biological replicates. Fold changes from qRT-PCR were compared with the RNA-seq expression profiles (***Figure 1H***). The dynamic expression patterns of all genes were consistent with the RNA-seq analysis, showing a high correlation (Pearson’s correlation coefficient of 0.892) between RNA-seq and qRT-PCR data. These results therefore indicated the reliability of the RNA-seq for mRNA differential expression analysis.

### Functional enrichment analysis of DEGs during spermatozoa differentiation and maturation

Gene ontology (GO) term-enriched analysis of DEGs in SPZ_EJ_ with significant differences revealed that a large number of biological processes were represented. The five top-ranked GO terms were regulation of biological, cellular and metabolic processes, and organic substance and metabolic processes (***Figure 2-Supplement 1A***). Further analysis of GO term distribution indicated that the most represented biological process was the regulation of gene expression, followed by positive regulation of macromolecule and cellular metabolism, regulation of signal transduction, and regulation of cellular biosynthesis (***Figure 2-Supplement 1B***). Interestingly, genes with GO terms such as cellular response to stimulus, cell communication, signal transduction, response to external or chemical stimulus, cell adhesion, and cell surface receptor signaling pathway, were only upregulated in SPZ_EJ_ (***Figure 2-Supplement 1A***). For the GO molecular function, the top enriched terms were binding to ribonucleotides and purine nucleotides, whereas the terms Ca^2+^, phosphatidylinositol and actin binding, ion channel activity, and transmembrane transport of inorganic cations and organic anions appeared to be only upregulated in SPZ_EJ_ (***Figure 2-Supplement 1C***). Taken together these findings indicate the enrichment of gene expression, metabolic and signaling processes in SPZ_EJ_.

To gather more information on genes with a potential impact on spermatozoa function, the DEGs in SPZ_EJ_ were manually classified into five functional categories by using GO analysis and the Uniprot database. These categories included transcription, translation and chromatin organization (1,056 genes), receptors (433 genes), metabolism of proteins, lipids and carbohydrates (492 genes), cytoskeleton and cell movement (520 genes), and channels, exchangers and transporters (308 genes) (***Figure 2A***). The genes upregulated in SPZ_EJ_ related to transcription, translation and chromatin organization (443 genes) mainly correspond to transcription factors (42.5%) and regulators of transcription (20.1%), followed by ribosome structure (12%), regulators of translation (5.4%), chromatin and RNA binding (4.1 and 4.7%, respectively), and histones and histone modification (6.8%) (***Figure 2B***). Most of the receptor-encoding upregulated genes (303 genes) were G protein-coupled receptors (36.6%), tyrosine phosphatase and kinase receptors (11.5%), cytokine receptors (7.3%), as well as other receptors mainly including Fc receptors and novel immune-type receptors (***Figure 2C***). For metabolic processes (253 genes), the most enriched genes in SPZ_EJ_ were those related to glycolysis and gluconeogenesis (8.7%), the metabolism of glycogen and other polysaccharides (11.8%), fatty acids (26.1%) and amino acids (11.5%), and proteases (17.4%) (***Figure 2D***). Finally, genes encoding for proteins involved in cytoskeletal organization (32.1%), actin binding (21.4%) and motor proteins (15.7%) were the most abundant upregulated genes involved in the cytoskeleton and cell movement (149 genes) (***Figure 2E***), whereas in the group including upregulated genes encoding for channels, exchangers and transporters (139 genes) the K^+^ and metal specific channels (19.9%), cation channels (13.2%) and peptide and amino acid transporters (16.9%) were the most enriched in SPZ_EJ_ (***Figure 2F***).

**Figure 2.**
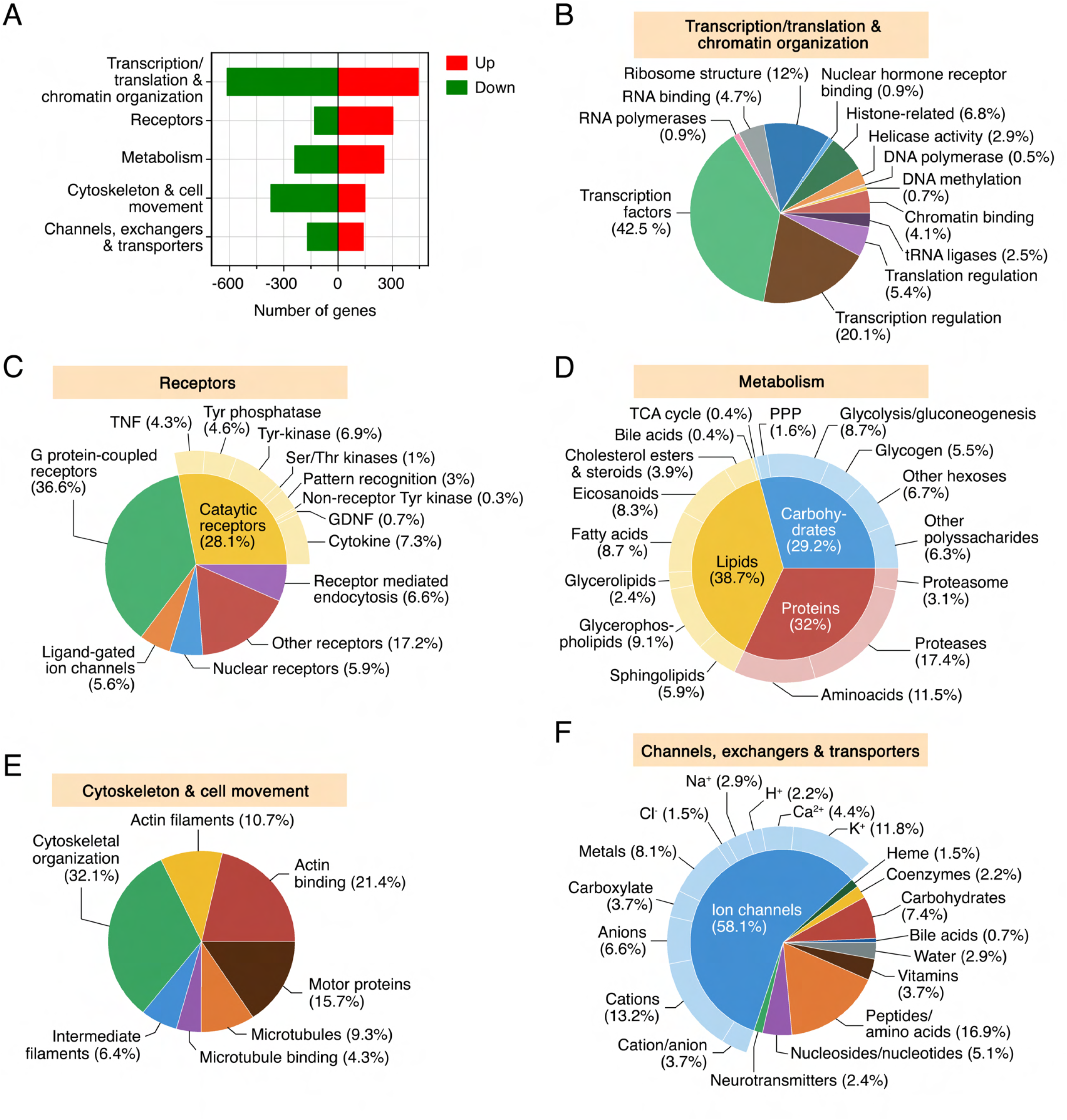
Functional classification of DEGs during sperm differentiation and maturation. (**A**) Transcriptional regulation of a subpopulation of DEGs classified into five functional categories: transcription and translation and chromatin organization, receptors, metabolism, cytoskeleton and cell movement, and channels, exchangers and transporters. (**B**-**F**) Pie charts showing the GO term distribution of upregulated DEGs in SPZ_EJ_ included in each of the five functional groups. The numbers are the percentage of genes in each category. **Figure 2-source data 1** Data for the classification of DEGs. **Figure 2-Supplement 1-source data 1** Data from GO analysis of the DEGs during sperm differentiation and maturation.

In an effort to identify specific transcription/translation and metabolic processes enriched in SPZ_EJ_, we built the protein interactome network of DEGs classified into these two categories by using the STRING protein-protein interaction (PPI) database for known PPIs (***Szklarczyk et al., 2019***) with very stringent inclusion criteria. As a result, a connected network comprising 766 proteins and 3,588 connections was mapped for the proteins encoded by genes involved in transcription and translation (***Figure 3A***). These proteins could be divided into five major subclusters based on their known biological functions established through GO analysis, including mitochondrial translation, tRNA aminoacylation, translation initiation, cytosolic ribosomes and mRNA splicing (***Figure 3A***). All of the DEGs grouped into the cytosolic ribosome subunit subcluster, and half of the DEGs belonging to the tRNA aminoacylation, mitochondrial translation, and translation initiation subclusters, were upregulated (***Figure 3A***). These findings, together with the previous observation of the high abundance of upregulated genes encoding for transcription factors and transcription regulators in SPZ_EJ_, suggest that both mitochondrial and cytoplasmic translation activity occurs during the differentiation and maturation of spermatozoa.

**Figure 3.**
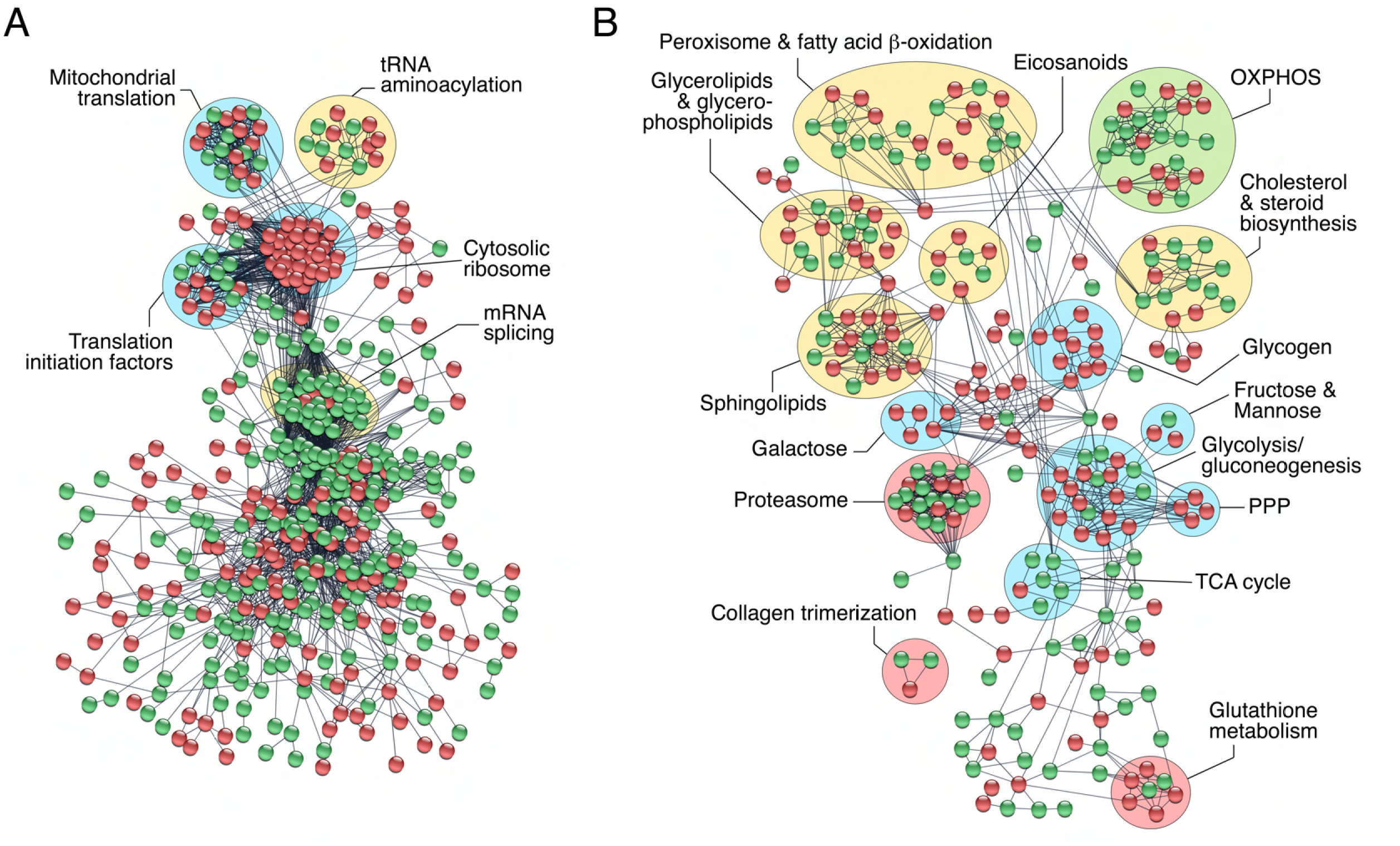
Protein-protein interaction (PPI) networks of DEGs. The PPI information of DEGs potentially involved in transcription and translation and chromatin organization (**A**), and metabolism (**B**), was obtained through a database search using STRING database v11 with a high confidence score (0.9), and imported into Cytoscape v3.8.2 for network construction. Proteins and their interactions are shown as nodes (spheres) and edges (lines), respectively. Nodes in red or green color indicate upregulated and downregulated DEGs, respectively. Proteins are grouped based on their known biological functions. Abbreviations: OXPHOS, oxidative phosphorylation; PPP, pentose phosphate pathway; TCA, tricarboxylic acid.

The metabolism interactome showed 379 proteins and 821 connections divided into fifteen subclusters, from which those corresponding to glycolysis/gluconeogenesis, pentose phosphate pathway (PPP) and sphingolipid, galactose, glycogen and glutathione metabolism, were the most upregulated in SPZ_EJ_ (***Figure 3B***). Further mapping of the 76 DEGs coding for enzymes involved in respiratory pathways indicated that most of the genes of the tricarboxylic acid (TCA) cycle, as well as three genes coding for specific enzymes of gluconeogenesis, such as phosphoenol-pyruvate carboxykinase (*pck2*), fructose 1,6-bisphosphatase (*fbp1*) and glucose 6-phosphatase (*g6pc*), were downregulated or not differentially expressed in SPZ_EJ_ (***Figure 3-Supplement 1***). In contrast, most of the glycolytic enzyme-encoding genes, including the two key enzymes hexokinase-1 (*hk1*) and pyruvate kinase (*pkm*), as well as many of the genes coding for enzymes catalyzing oxidative phosphorylation (OXPHOS), were upregulated (***Figure 3-Supplement 1***), suggesting that both glycolysis and OXPHOS are possibly important pathways for ATP generation in seabream spermatozoa.

### The PDGF and GnRHR signaling pathways are upregulated in SPZ_EJ_

In order to identify signaling pathways enriched in SPZ_EJ_, pathway analysis was done for the 7,287 DEGs using the PANTHER classification system (***Mi et al., 2019***). The analysis identified a total of 960 transcripts belonging to 37 different signaling pathways, including 13 receptor pathways (***Figure 4A***). Highly enriched and significant pathways were integrin, epidermal growth factor receptor (EGFR), fibroblast growth factor (FGF), cholecystokinin receptor (CCKR) and inflammation mediated by chemokine and cytokine. These results possibly reflect the activation of mechanisms for actin remodeling (Breitbart et al., 2011), the acquisition and regulation of motility and chemotaxis (***Caballero-Campo et al., 2014***; ***Tan and Thomas, 2015***; ***Zhou et al., 2015***; ***Saucedo et al., 2018***), and the formation of an active network of proteins prior to fertilization crucial for the sperm-egg fusion (***Chen et al., 1999***; ***Frolikova et al., 2016***), during the differentiation and maturation of spermatozoa.

**Figure 4.**
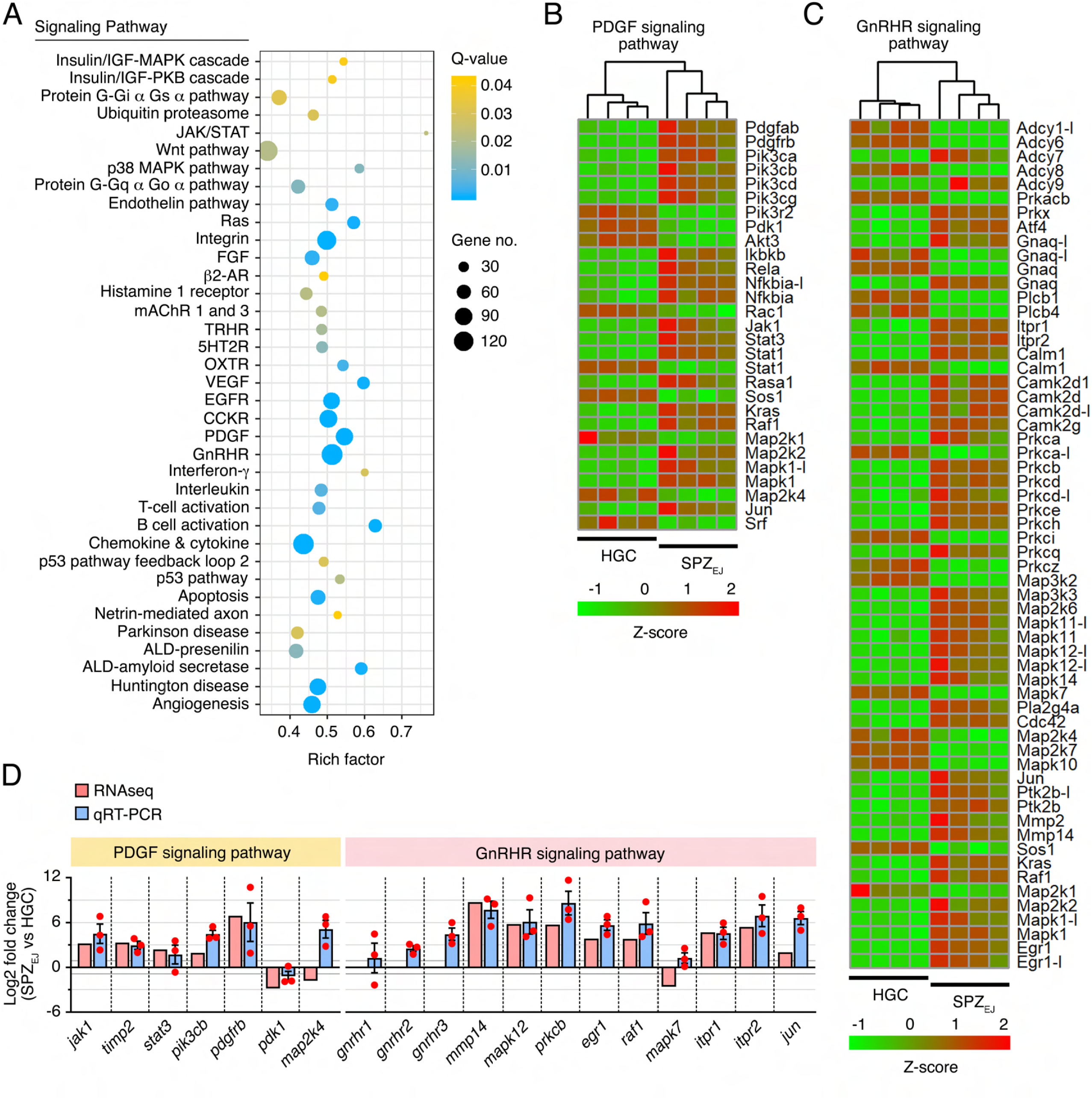
Pathway enrichment analysis during spermatozoa differentiation and maturation. **(A)** Pathway analysis of DEGs using the PANTHER Classification System showing the 37 most highly enriched signaling pathways (FDR < 0.05) in SPZ_EJ_. (**B** and **C**) Hierarchical clustering heatmaps of DEGs related to the PDGF (B) and GnRHR (C) signaling pathways. (**D**) qRT-PCR validation of the changes in expression of several genes classified into the PDGF or GnRHR pathways. Data from qRT-PCR are the mean ± SEM (*n* = 3 pools of 3 different fish each). **Figure 4-source data 1** Data for heatmap shown in B. **Figure 4-source data 2** Data for heatmap shown in C. **Figure 4-source data 3** Data on the qRT-PCR validation of the changes in expression of several genes classified into the PDGF or GnRHR pathways.

However, amongst the most dominant pathways in terms of number of genes identified and lowest *p*-values were the GnRHR and PDGF signaling pathways. Hierarchical clustering heatmaps showed that most of the genes related to these two pathways encoding for receptors, kinases, transcription factors or calcium binding proteins were upregulated, such as the Pdgf receptor b (*pdgfrb*), phosphatidylinositol 4,5-bisphosphate 3-kinase (*pik3*), nuclear factor kappaB subunit p65 (*rela*), NF-kappa-B inhibitor alpha (*nfkbia*), signal transducer and activator of transcription 1 (*stat1*), GTPase Kras (*kras*), RAF proto-oncogene Ser/Thr-protein kinase (*raf1*), mitogen-activated protein kinase 1 (*mapk1*), dual specificity mitogen-activated protein kinase kinase 2 (*map2k2*) or transcription factor AP-1 (*jun*) in the PDGF pathway (***Figure 4B***), and adenylate cyclase type 7 (*adcy7*), cAMP-dependent protein kinase catalytic subunit PRKX (*prkx*), c-AMP-dependent transcription factor ATF-4 (*atf4*), guanine nucleotide-binding protein G(q) subunit alpha (*gnaq*), inositol 1,4,5-trisphosphate receptor type 1 and 2 (*itpr1* and *itpr2*), calmodulin-1 (*calm1*), protein kinase beta, delta and epsilon (*prkcb*, *prkcd* and *prkce*) or early growth response protein 1 (*egr1*) (***Figure 4C***), in the GnRHR pathway. These data were validated by qRT-PCR for a number of genes, including three Gnrhrs identified in our transcriptome (*gnrhr1*, *gnrhr2* and *gnrhr3*) for which the RNA-seq did not detect significantly different expression levels (***Figure 4D***). The qRT-PCR analysis showed however that *gnrhr2* and *gnrhr3* were in fact upregulated in SPZ_EJ_ (***Figure 4D***) Altogether, these data suggest the activation of the GnRHR and PDGF signaling pathways during seabream spermiogenesis.

### Seabream Gnrh and Pdgf paralogs are sequentially expressed in the ETD epithelia

Since seabream SPZ_EJ_ show elevated expression of components of the GnRHR and PDGF signaling pathways, we speculated that these pathways might be involved in the maturation of the spermatozoon in the ETDs, the efferent (ED) and sperm (SD) ducts (Figure 5*A* and *B*), prior to ejaculation. To investigate this hypothesis, we first evaluated whether two molecular forms of Gnrh present in gilthead seabream, the seabream Gnrh (*sbgnrh*) and salmon Gnrh (*sgnrh*) (***Powell et al., 1994***), as well as different Pdgf paralogs identified in our transcriptome and in the seabream genome (*pdgfaa*, *-ab*, *-ba*, *-bb*, *-c* and *-d*), are expressed in the testis, ED and SD. *In situ* hybridization using DIG-labeled, paralog-specific riboprobes showed strong *sbgnrh* expression in SPC in the testis, whereas the expression was also prominent in nonciliated cells lining the lumen of the ED (***Figure 5C***). In contrast, *sbgnrh* transcripts were almost undetectable in the ciliated epithelial cells of the proximal and distal regions of the SD (***Figure 5C***). The *sgnrh* mRNA was also detected exclusively in testicular SPC, while a faint signal was observed in the epithelial cells from ED but not from the SD (***Figure 5-Supplement 1***). The expression of the *sbgnrh* paralog correlated with the immunostaining of Gnrh peptides using an anti-GnRH antibody, confirming that the sbGnrh was produced in testicular SPC and nonciliated epithelial cells of the ED, the expression of the neuropeptide being progressively decreased along the SD (***Figure 5D***).

**Figure 5.**
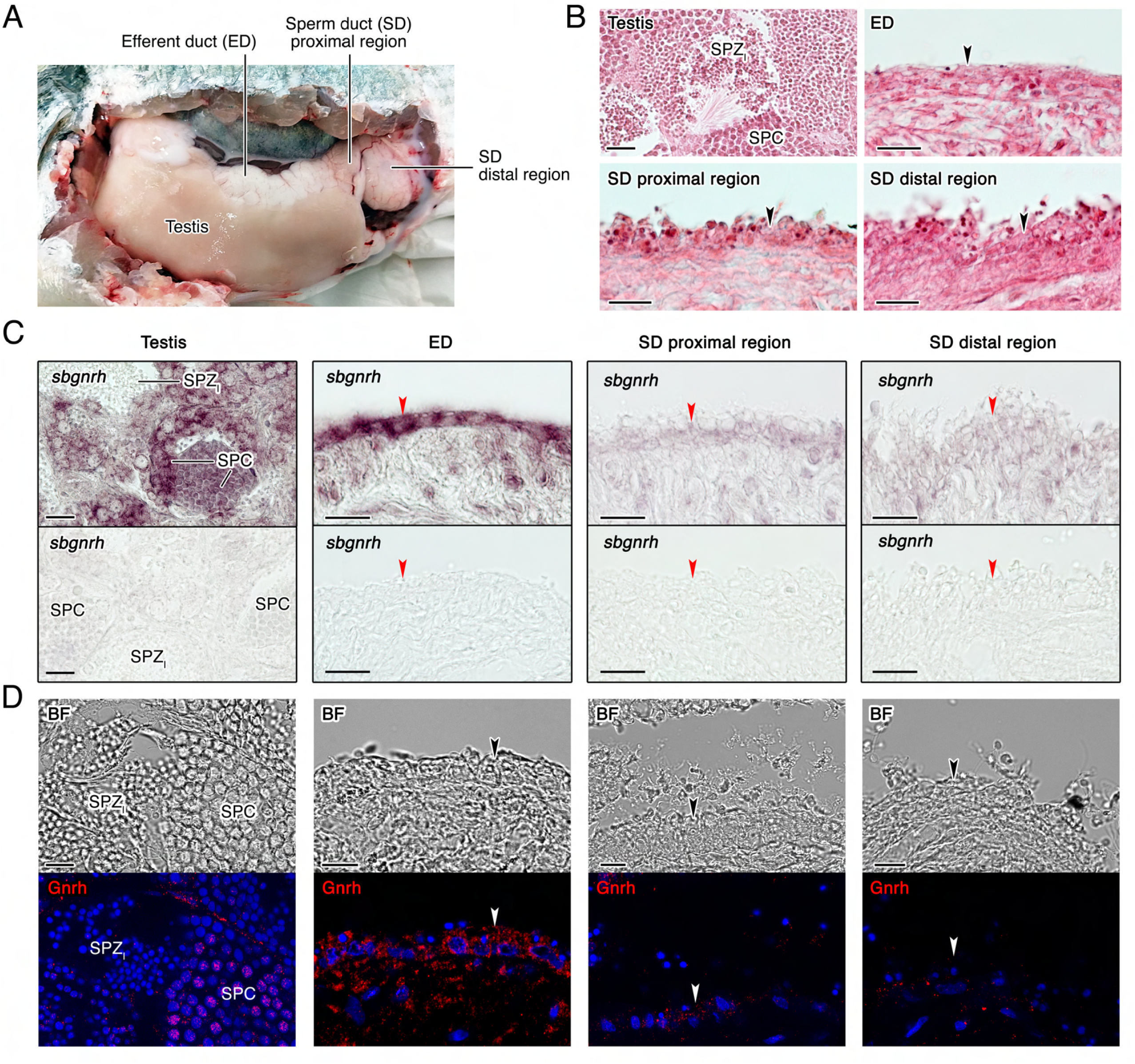
Cellular localization of GnRH expression in seabream extratesticular ducts. (**A**) Anatomy of the seabream testis and extratesticular ducts, efferent duct (ED) and sperm duct (SD). (**B**) Paraffin histological sections of the different structures of the testis and testicular ducts stained with hematoxylin and eosin. (**C**) Localization of *sbgnrh* transcripts in the testis, ED and SD by *in situ hybridization* on paraffin sections hybridized with antisense DIG-labeled riboprobes specific for *sbgnrh* (upper panels) or sense probes (lower panels, negative controls). (**D**) Immunostaining of GnRH peptides (red, lower panels) in the same testicular structures as in C. Corresponding brightfield (BF) images are also shown (upper panels). The reactions were visualized with Cy3-conjugated sheep anti-rabbit IgG and the nuclei were counterstained with 4’,6-diamidino-2-phenylindole (DAPI; blue). Control sections incubated with the secondary antibody only did not show any staining (**Figure 5-*Supplement 2***). Scale bars, 50 µm (B and C), 10 µm (D). Abbreviations: SPC, spermatocytes; SPZ_I_, intratesticular spermatozoa. The arrowheads in B-D indicate epithelial cells of the ED and SD.

The cell localization of *pdgf* expression in testis and ETDs by *in situ* hybridization revealed distinct expression patterns of the different *pdgf* paralogs. In the testis, expression of *pdgfaa* was specific of the somatic Sertoli cells, but only when they showed embedded developing spermatogonia (SPG) (***Figure 6-Supplement 1***), while no positive signals were detected for *pdgfab* (***Figure 6A***). The *pdgfba* mRNA was detected in SPG, with much weaker signals in SPC and SPD (***Figure 6B***), whereas the transcripts of the duplicated *pdgfbb* paralog, as well as those of the *pdgfd*, were also localized in SPC but they were much less abundant in SPD (***Figure 6C and D***). The expression of *pdgfc* was only detected in Leydig cells (***Figure 6-Supplement 1***). In the ETDs, *pdgfab* expression was strong in the epithelial cells of the ED, being weak in the SD (***Figure 6A***), whereas the expression of *pdgfba* and *-bb* was more intense in the luminal surface of the SD proximal and distal regions, respectively (***Figure 6B and C***). In contrast, *pdgfd* expression was low in the epithelium throughout the ETDs, but somewhat more intense in the proximal region of the SD (***Figure 6D***), while the expression of *pdgfaa* and *-c* was almost or completely undetectable (***Figure 6-Supplement 1***).

**Figure 6.**
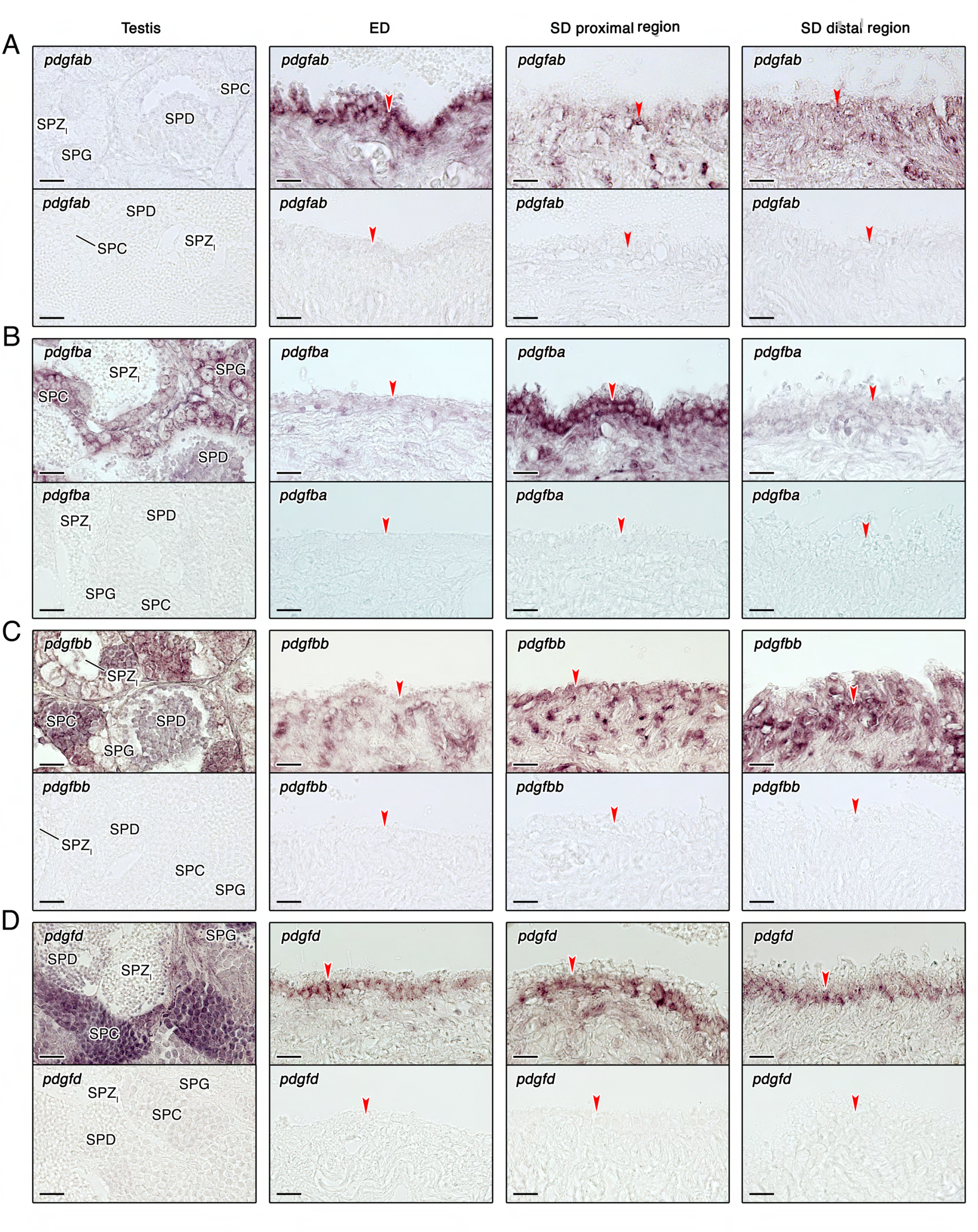
Localization of *pdgf* transcripts in the seabream testis, ED and SD. (**A**-**D**) Paraffin sections were hybridized with antisense DIG-labeled riboprobes specific for different *pdgf* paralogs (upper panels) as indicated. Control sections (lower panels), hybridized with sense probes, were negative. Scale bars, 50 µm. SPG, spermatogonia; SPC, spermatocytes; SPD, spermatids; SPZ_I_, intratesticular spermatozoa. The arrowheads indicate epithelial cells of the ED and SD.

Taken together, these findings demonstrate the local production of Gnrh peptides by the epithelial cells of the ED, as well as the sequential expression of different *pdgf* paralogs along the ETDs, which would be consistent with a physiological role of these hormones during the maturation of spermatozoa.

### GnRH and PDGF regulate transcription and motility of immature seabream spermatozoa

To investigate the physiological state of the spermatozoa from the ED (SPZ_ED_), we compared their function with respect to that of SPZ_EJ_. Time-course monitoring of sperm motion kinetics upon seawater activation using computer-assisted sperm analysis (CASA) revealed that SPZ_ED_ showed a reduced percentage of motility and progressivity, and an impaired curvilinear velocity (VCL), with respect the SPZ_EJ_ (***Figure 7A***). These data therefore indicate that SPZ_ED_ can be classified as immature gametes, which acquire full motility potential during their journey throughout the ETDs.

**Figure 7.**
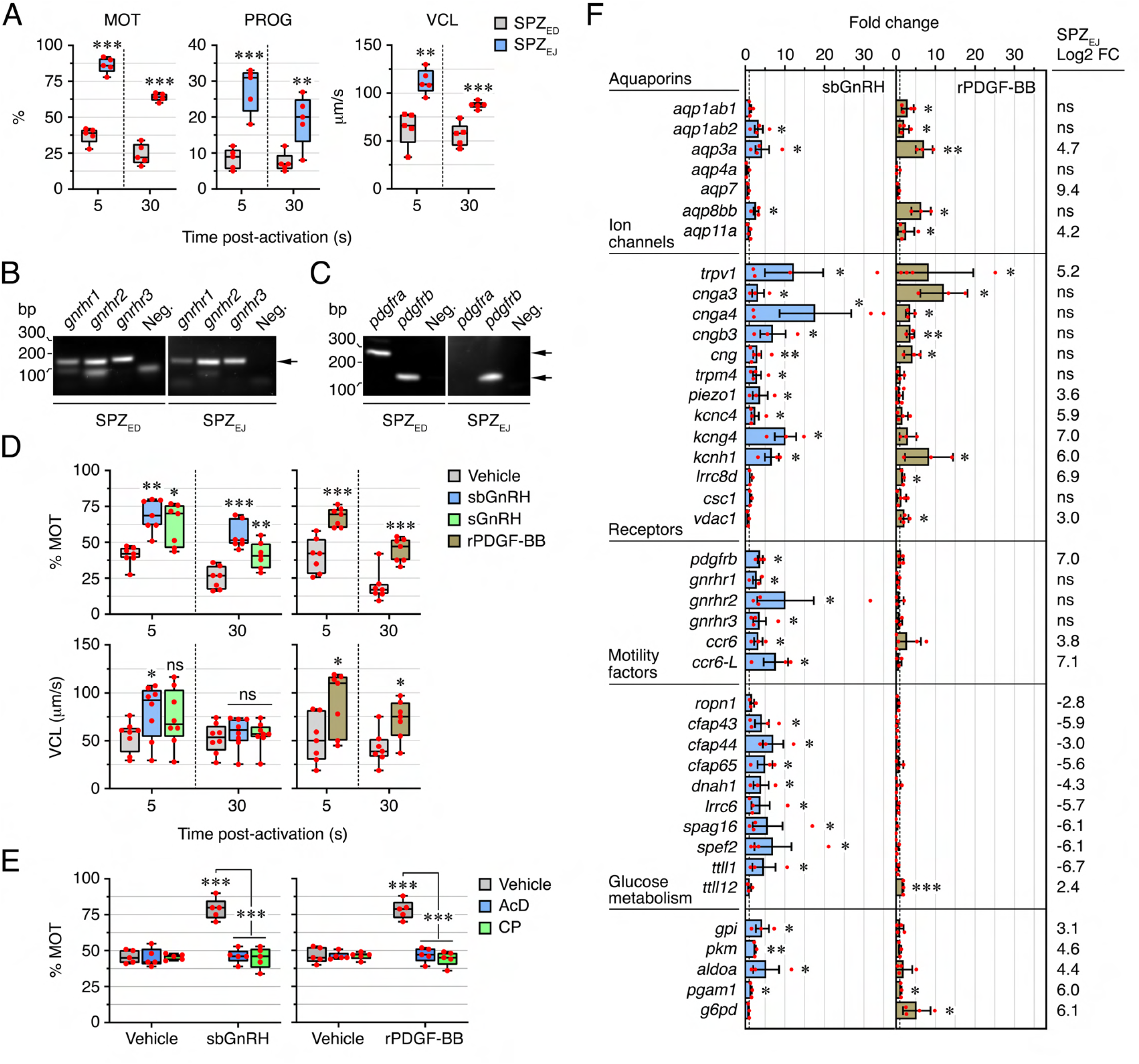
Transcriptional regulation of seabream sperm motility by GnRH and PDGF. (**A**) Percentage of motility (%MOT) and progressivity (%PROG), and curvilinear velocity (VCL), of spermatozoa from the efferent duct (SPZ_ED_) or ejaculated (SPZ_EJ_) determined at 5 or 30 s post activation. (**B** and **C**) RT-PCR detection of mRNAs encoding GnRH receptors (*gnrhr1*, *gnrhr2* and *gnrhr3*) and PDGF receptors b (*pdgfra* and *pdgfrb*) in SPZ_ED_ or SPZ_EJ_. The Neg. line is the negative control (absence of RT during cDNA synthesis). The arrows indicate the specific transcripts, and the size (kb) of molecular markers are indicated on the left. (**D**) The % MOT and VCL of SPZ_ED_ exposed to 100 nM of sbGnRH or sGnRH, 40 nM of mouse recombinant PDGF (rPDGF-BB), or to each hormone vehicle, determined at 5 or 30 s post activation. (**E**) Inhibition of motility of SPZ_ED_ induced by sbGnRH and rPDGF-BB by 100 µg/ml actinomycin D (AcD) or chloramphenicol (CP) at 5 s post activation. (**F**) Quantitative RT-PCR analysis of the expression of selected genes potentially involved in water and ion transport, signaling, flagellar motility and glucose metabolism in SPZ, after sbGnRH or rPDGF-BB stimulation. The Log2 fold change in the expression of each gene in the RNA-seq analysis is indicated on the right. In A, D and E, all data points are presented as box and whisker plots/scatter dots with horizontal line (inside box) indicating median and outliers. One ejaculate from each male was measured from *n* = 5-7 males. In F, data are the mean ± SEM (*n* = 3-5 fish). Data were statistically analyzed by an unpaired Student’s *t*-test (A and F), or by one-way ANOVA (D and E). *, *P* < 0.05; **, *P* < 0.01; ***, *P* < 0.001; with respect to spermatozoa incubated with the hormone vehicles, or as indicated in brackets. **Figure 7-source data 1** Data on sperm motility shown in A. **Figure 7-source data 2** Uncropped gels of the RT-PCR of mRNAs encoding seabream GnRH receptors (*gnrhr1*, *gnrhr2* and *gnrhr3*) in SPZ_ED_ or SPZ_EJ_. The Neg. line is the negative control (absence of RT during cDNA synthesis). The arrows indicate the specific transcripts, and molecular markers are on the left. **Figure 7-source data 3** RT-PCR detection of mRNAs encoding seabream PDGF receptors b (*pdgfra* and *pdgfrb*) in SPZ_ED_ or SPZ_EJ_. The Neg. line is the negative control (absence of RT during cDNA synthesis). The arrows indicate the specific transcripts, and the molecular markers are on the left. **Figure 7-source data 4** Data on sperm motility shown in D. **Figure 7-source data 5** Data on sperm motility shown in E **Figure 7-Suplement 1-source data 1** Data on sperm motility shown in Figure 7-Supplement 1. **Figure 7-source data 6** Quantitative RT-PCR analysis of the expression of selected genes shown in F.

Further RT-PCR analysis showed that both SPZ_ED_ and SPZ_EJ_ express *gnrhr1*, *gnrhr2* and *gnrhr3* transcripts (***Figure 7B***), while expression of *pdgfra* is specific of SPZ_ED_, and that of *pdgfrb* is prevalent in SPZ_ED_ and SPZ_EJ_ (***Figure 7C***). Therefore, we tested the hypothesis that the activation of these receptors by their cognate ligands in SPZ_ED_ may play a role in the acquisition of full motility. For this, SPZ_ED_ were incubated with sbGnRH, sGnRH or mouse recombinant PDGF-BB (rPDGF-BB), and subsequently activated in seawater to determine changes in motility. Exposure to the three hormones significantly increased the motility, progressivity and VCL of the SPZ_ED_, although the positive effect of rPDGF-BB on the velocity appeared to be more persistent over time than that of sbGnRH (***Figure 7D and Figure 7-Supplement 1***). However, the stimulation of SPZ_ED_ motility by both sbGnRH and rPDGF-BB was completely abolished by preincubation of spermatozoa with the transcription inhibitor actinomycin D or the mitochondrial translation inhibitor chloramphenicol (***Figure 7E and Figure 7-Supplement 1***), suggesting that the sbGnRH- and rPDGF-BB-mediated regulation of motility is dependent on transcription and mitochondrial translation in spermatozoa.

To investigate the transcription-dependent actions of sbGnRH and PDGF on the motility of SPZ_ED_, we employed a targeted approach by evaluating the hormone-induced changes in the expression levels of selected genes. This included genes that encode aquaporins and ion channels, receptors, components of the motile apparatus, and enzymes involved in respiratory pathways, most of them regulated in SPZ_EJ_ as indicated by RNA-seq profiling (***Figure 2 and 3***), and which control or can potentially modulate sperm motility (***Boj et al., 2015***; ***Chauvigné et al., 2015***; ***Alavi et al., 2019***; ***Chauvigné et al., 2021***). The data indicated that both sbGnRH and rPDGF-BB upregulated the expression of many of these genes in SPZ_ED_, but not all the same genes were affected by the two hormones (***Figure 7F***). Thus, sbGnRH stimulated the expression of *aqp1ab2*, -*3a* and -*8bb*, whereas rPDGF-BB increased the amount of the same transcripts as well as those of *aqp1ab1* and -*11a*. In contrast, sbGnRH induced higher expression levels of different ion channels (*trpv1*, *cnga3*, *-4*, *cngb3*, *cng*, *trpm4*, *piezo1*, *kcnc4*, *kcng4*, and *kcnh1*) compared to rPDGF-BB, whereas the growth factor upregulated some of the same genes (*trpv1*, *cnga3*, *-4*, *cngb3*, and *cng*), as well as that of *lrrc8d* and *vdac1*, which were not regulated by the neuropeptide. Interestingly, all the receptors analyzed (*gnrhr1*, *gnrhr2*, *gnrhr3*, *pdgfrb*, *ccr6* and *ccr6-L*) and most of the genes related to sperm flagellar motility (*cfap43*, *-44*, *-65*, *dnah1*, *lrrc6*, *spag16*, *spef2*, and *ttll1*) were upregulated only by sbGnRH, while the rPDGF-BB exclusively increased the expression of *ttll12*. Finally, the data showed that sbGnRH stimulated the expression of several glycolytic enzymes, such as *gpi*, *pkm*, *aldoa* and *pgam1*, whereas rPDGF-BB only upregulated *pgam1* and *g6pd*, the latter enzyme catalyzing the rate-limiting step of the PPP. Many of the genes upregulated by sbGnRH or rPDGF-BB in SPZ_ED_ *in vitro* also appeared to be upregulated in SPZ_EJ_ *in vivo* in the RNA-seq (***Figure 7F***). However, most of the sbGnRH regulated genes coding for motility factors were downregulated in SPZ_EJ_ (***Figure 7F***), which may reflect an early and transitory activation of these genes during the maturation of spermatozoa in the ED *in vivo*. In any case, our findings suggest that both sbGnRH and PDGF play a role in the maturation of SPZ_ED_ through transcriptional activation of genes involved in the acquisition and maintenance of flagellar motility.

### Transcription-dependent regulation of sperm maturation by GnRH and PDGF is conserved in zebrafish

To examine whether the GnRHR and PDGF signaling pathways regulating seabream sperm maturation could be conserved in teleosts from more ancestral lineages, we first localized both GnRH and PDGF expressing cells in the testis and ETDs of the zebrafish (***Figure 8A***). Immunostaining and *in situ* hybridization experiments showed the expression of GnRH peptides, as well as of *pdgfaa* and *-bb* transcripts, in the epithelial cells lining the ETD (***Figure 8B-D***), thus suggesting the existence of the epithelial GnRH and PDGF signaling pathways in the ETD of zebrafish as observed in seabream.

**Figure 8.**
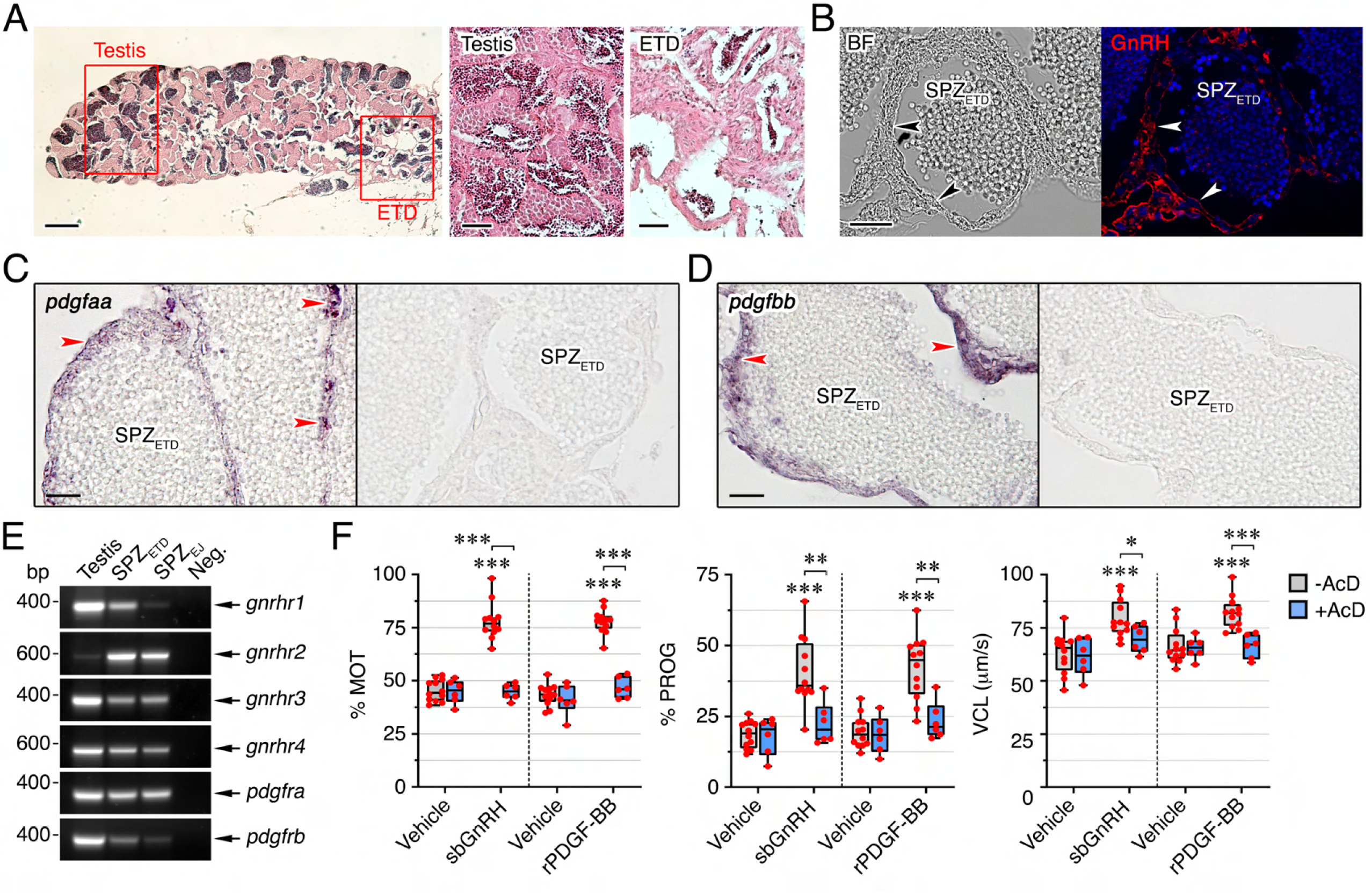
Transcription-dependent regulation of zebrafish sperm motility by GnRH and PDGF. (**A**) Paraffin histological sections of the zebrafish testis and extratesticular ducts (ETD) stained with hematoxylin and eosin. Scale bars, 10 and 100 µm. (**B**) Immunostaining of GnRH peptides (red, right panel) in the surface epithelium of the ETDs (arrowheads) and corresponding brightfield (BF) image (left panel). Control sections incubated with the secondary antibody only were negative (**Figure 8-*Supplement 1***). Scale bar, 200 µm. (**C** and **D**) Paraffin sections the ETDs hybridized with antisense DIG-labeled riboprobes specific for *pdgfaa* (C) and *pdgfbb* (D) mRNAs as indicated. The arrowheads indicate expression in the epithelial cells of the ETDs. The right panels show the absence of signals in sections hybridized with sense probes. SPZ_ETD_, spermatozoa from the ETDs. (**E**) RT-PCR detection of mRNAs encoding GnRH (*gnrhr1*, *gnrhr2*, *gnrhr3* and *gnrhr4*) and PDGF (*pdgfra* and *pdgfrb*) receptors in SPZ_ETD_ and SPZ_EJ_. The Neg. line is the negative control (absence of RT during cDNA synthesis). The arrows indicate the specific transcripts, and the size (kb) of molecular markers are indicated on the left. (**F**) Percentage of motility (%MOT) and progressivity (%PROG), and curvilinear velocity (VCL), at 5 s postactivation of SPZ_ETD_ exposed to 100 nM of sbGnRH, 40 nM of mouse recombinant PDGF (rPDGF-BB), or to each hormone vehicle, in the presence or absence of actinomycin D (AcD; 100 µg/ml). Data are presented as box and whisker plots/scatter dots with horizontal line (inside box) indicating median and outliers (*n* = 6-12 fish), and were statistically analyzed by an unpaired Student’s *t*-test. *, *P* < 0.05; **, *P* < 0.01; ***, *P* < 0.001; with respect to spermatozoa incubated with the hormone vehicles, or as indicated in brackets. **Figure 8-Supplement 2-source data 1** Data on sperm kinetics shown in **Figure 8-**Supplement 2. **Figure 8-source data 1** Uncropped gels from RT-PCR detection of mRNAs encoding zebrafish GnRH (*gnrhr1*, *gnrhr2*, *gnrhr3* and *gnrhr4*) and PDGF (*pdgfra* and *pdgfrb*) receptors in SPZ_ETD_ and SPZ_EJ_. The Neg. line is the negative control (absence of RT during cDNA synthesis). The arrows indicate the specific transcripts, and the molecular markers are on the left. **Figure 8-source data 2** Data on sperm motility shown in F.

To further assess whether sbGnRH and rPDGF-BB can induce sperm maturation in zebrafish, we first confirmed that spermatozoa from the ETD (SPZ_ETD_) show lower motility than SPZ_EJ_ upon activation in freshwater (***Figure 8-Supplement 2***), and that they express the four GnRH (*gnrhr1*, *-r2*, *-r3* and *-r4*) and two PDGF (*pdgfra* and *-b*) receptors formerly identified in zebrafish (***Tello et al., 2008***; ***Eberhart et al., 2008***) (***Figure 8E***). This allowed us to classify the zebrafish SPZ_ETD_ as immature sperm cells. *In vitro* incubation of SPZ_ETD_ with sbGnRH or rPDGF-BB before activation significantly increased the motility, progressivity and VCL of the spermatozoa, each of which was completely blocked by the addition of actinomycin D (***Figure 8F***). These data demonstrate that the transcription-dependent maturation of SPZ_ETD_ induced by GnRH and PDGF is a conserved mechanism in teleosts.

## Discussion

The present results reveal that the widely accepted view of virtual transcriptional silence during post-meiotic spermatozoon maturation (Fisher et al., 2012; ***Grunewald et al., 2005***; ***Ren et al., 2017***; ***Freitas et al., 2020***; Puga Molina et al., 2018) is not a conserved phenomenon in vertebrates. By selecting a species of fish that does not incorporate protamines during the chromatin condensation phase of spermiogenesis and conducting transcriptome profiling and gene set enrichment analysis during spermatozoon differentiation, we uncovered novel endocrine signaling pathways in the ETD required for the acquisition of sperm motility competence. This mechanism does not rely on the delivery of specific mRNAs via epididysome-like vesicles as in mammals (***James et al., 2020***), but on *de novo* transcription and translation mechanisms occurring in maturing spermatozoa.

In most teleosts, sperm maturation, the phase during which non-functional gametes develop into mature spermatozoa, fully capable of vigorous motility and fertilization, is believed to occur in the ETDs (***Schulz et al., 2010***). Previous studies have shown that administration of some hormones, such as progestins, androgens and gonadotropins, can increase the seminal plasma pH in the ETD, which results in the elevation of intra-sperm cAMP levels, increase hydration, or induce the secretion of sperm-immobilizing ions by the ETD epithelium (***Schulz et al., 2010***; ***Marshall et al., 1993***). However, the cellular sources of these hormones in the ETD and their potential signal transducing effects in the maturing spermatozoa are completely unknown. In the present study, the transcriptomic analysis of the enriched signaling pathways in seabream SPZ_EJ_ revealed the presence of conserved GnRH and PDGF endocrine pathways in the teleost ETD. These hormones are produced in a spatially distinct expression sequence by the epithelial cells lining the ED and SD in a manner that resembles the regional expression of genes in the mammalian epididymis (***Sullivan and Mieusset, 2016***; ***Belleannée et al., 2012***; ***Zhao et al., 2019***). Such spatiotemporal expression is thus likely an ancient signaling mechanism that evolved early in the development of ETDs in jawed vertebrates. In the present context, therefore, different paralogs of the GnRH and PDGF hormones provide a programmed sequence of paracrine signals to activate their cognate receptors in SPZ_ED_ to transduce *de novo* transcription and translation.

To validate this model, we performed *in vitro* experiments to maturationally induce SPZ_ED_ with sbGnRH or rPDGF-BB, and conducted qRT-PCR of 45 different genes, which regulate or can potentially modulate sperm motility. We further validated the maturational status of the endocrine-induced and non-induced SPZ_ED_ in the presence and absence of the transcription inhibitor actinomycin D following seawater activation and CASA analysis. These data show that for both seabream and zebrafish, the motility of the SPZ_ED_ only increases in the presence of hormones, and only in the absence of the transcriptional inhibitor. For seabream these experiments also demonstrate the importance of mitochondrial translation for the increase in motility, as observed in mammals (***Gur and Breitbart, 2006***; ***Zhao et al., 2009***; ***Rajamanickam et al., 2017***; ***Zhu et al., 2019***), while the qRT-PCR data confirm the effect of the hormones on the upregulation of a suite of downstream effector genes. In this latter respect, sbGnRH and rPDGF-BB show similar regulatory induction of aquaporins and ion channels, but sbGnRH has a more potent effect on the *de novo* transcription of receptors, motility factors and some key enzymes in glucose metabolism. Since we show that the sbGnRH neuropeptide is primarily expressed in the ED epithelium, the data suggest that the upregulation of sperm receptors, motility factors and genes associated with glucose metabolism is induced early in the maturational process. Conversely, the sequential expression of paralogous seabream *pdgf* receptors in separate regions of the ED and SD, indicate that regulation of aquaporins and ion channels occurs throughout the ETD.

The early upregulation of *pdgfrb* and *gnrhr1, -2 and -3* by sbGnRH in SPZ_ED_ is a clear indication of the acquisition of developmental competence, since expression of these receptors in the maturing spermatozoa assembles the signal transduction pathways capable of responding to the cognate hormones that are secreted from the somatic ETD. Thus, although region-specific gene expression is known in the mammalian epididymis (***Sullivan and Mieusset, 2016***; ***Belleannée et al., 2012***; ***Zhao et al., 2019***), and GnRH receptors are known to be expressed in primate spermatozoa (***de Villiers et al., 2021***), to the best of our knowledge, the epithelial ETD endocrine transduction of receptor-mediated gene transcription of the maturing spermatozoa has not previously been reported for vertebrates.

Interestingly, several of the genes that are hormonally upregulated in the maturing seabream spermatozoa have been shown to play important roles during the activation and maintenance of sperm motility. This includes aquaporins that facilitate the efflux of water for motility activation (***Boj et al., 2015***; ***Chauvigné et al., 2013***), or the mitochondrial efflux of hydrogen peroxide for the maintenance of ATP production and flagellar contractions (***Chauvigné et al., 2015***, ***2021***). Other upregulated genes encode ion channels involved in sperm motility, such as Trpv1 (***Majhi et al., 2013***; ***Chen et al., 2020***) and different cyclic nucleotide-gated channels (***Fechner et al., 2015***), or which are potentially implicated in cell volume regulation, such as Vdac1 (***Triphan et al., 2008***), Lrrc8d (***Jentsch, 2016***) and Trpv4 (***Benfenati et al., 2011***). Each of these two processes is considered important for the activation and maintenance of sperm motility in marine teleosts (***Boj et al., 2015***; ***Alavi et al., 2019***). The motility factors upregulated by sbGnRH were flagellar proteins involved in sperm flagellum axoneme organization and function (Cfap, Spag16, Spef2, Ttll1 and Ttll12) and dynein motor proteins (Dnah1, Lrcc6), which are likely necessary for flagellar function in piscine spermatozoa as in mammals (***Vogel et al., 2010***, ***Sironen et al., 2011***; ***Dzyuba and Cosson, 2014***; Feng et al., 2020; ***Wu et al., 2021***). The sbGnRH also activated the expression of Ropn1 early in the maturation process, which is an axonemal protein that plays a role in PKA-dependent signaling cascades required for spermatozoon capacitation (Fiedler at al., 2013). These findings therefore reinforce the notion that GnRH and PDGF signaling from the ETD epithelium plays a paracrine role to specifically induce the maturational expression of genes required for the activation and prolongation of sperm motility in the external aquatic environment.

In mammals, the vast majority of the paternal genome is packaged in protamines with transcriptional silence being coupled to heterochromatin condensation (***Sassone-Corsi, 2002***). In anamniotes, however, protamines may not be involved in nuclear chromatin condensation, or are completely lacking from the spermatozoon nucleus as in the species selected in the present study (***Shimizu et al., 2000***; ***Kurtz et al., 2009***; ***Wu et al., 2011***; ***Wike et al., 2021***). In seabream and zebrafish, spermiogenic nuclear chromatin condensation occurs without the replacement of the somatic-like and H1-family linker histones, so that they retain the nucleosome organisation with their nuclei remaining less condensed than those of species that incorporate protamines (***Kurtz et al., 2009***; ***Wike et al., 2021***; ***Saperas et al., 1993***). This is due to the absence of a second phase of spermiogenic chromatin condensation, which occurs when histones are displaced by SNBPs or protamines (***Kurtz et al., 2009***; ***Saperas et al., 1993***). In such cases, and indeed in a highly diverse range of species, including invertebrates, the first chromatin condensation transition is also characterized by low level acetylation that is not related to histone replacement (***Kurtz et al., 2007***, ***2009***). It seems plausible that the *de novo* transcription observed for the maturing spermatozoa of seabream in the present study, may therefore occur during this phase. In any event, the regional signaling of the ETD appears to be conserved in the epididymis of amniotes, but not the spermatozoon transcription. Future studies should investigate the chromatin architecture reorganization and epigenetic marks in teleost SPZ_ED_ that allow transcription and translation at this stage.

In summary, using a combination of transcriptional profiling, immunolocalization, *in situ* hydridization, and *in vitro* induction and inhibition experiments of sperm motility, we uncover novel endocrine signaling pathways in the ETD epithelium that transduce the *de novo* transcription of gametic effector genes required for fish sperm maturation. The experiments confirmed that the requirement of mitochondrial translation for the acquisition of full sperm motility is conserved between amniotes and anamniotes, but that transcriptional silence of post-meiotic spermatozoa is not a pan vertebrate phenomenon. In fishes, *de novo* transcriptional activation induced by soma to gamete signal transduction pathways is necessary for the acquisition of fertility competence.

## Materials and Methods

### Animals and sample collection

Adult gilthead seabream males were raised in captivity at Institut de Recerca i Tecnologia Agroalimentàries (IRTA) aquaculture facilities in San Carlos de la Rápita (Tarragona, Spain) and maintained in the laboratory as previously described (***Chauvigné et al., 2013***). Samples of testis and SPZ_EJ_ were obtained from males during the natural reproductive season (November-February) as previously described (***Chauvigné et al., 2013***), whereas the SPZ_ED_ was extracted with a micropipette after an incision in the dorsal region of the dissected testis close to the SD. Zebrafish were obtained from the PRBB Animal Facility (Barcelona, Spain) and kept at 28°C with 14-hour light and 10-hour dark cycle and fed daily with dry small granular pellets (Zebrafish Management Ltd) and newly hatched brine shrimp *Artemia franciscana*. To collect SPZ_EJ_, males were anaesthetised with 100 ppm 2-phenoxyethanol and euthanized, and the testis mixed with 30 µl of non-activating SS300 solution (in mg/ml: 8.15 NaCl, 0.67 KCl, 0.11 CaCl2, 0.12 MgSO4, 0.18 glucose, 2.42 Tris-Cl pH 8.0; 300 mOsm) (***Chauvigné et al., 2021***). Subsequently, the testis was mixed with 30 µl of fresh SS300 solution and slightly crushed using a micropippete to isolate SPZ_ETD_.

Procedures relating to the care and use of animals and sample collection were approved by the Ethics Committee (EC) of Institut de Recerca i Tecnologia Agroalimentàries (IRTA) and Universitat Autònoma de Barcelona (UAB), following the International Guiding Principles for Research Involving Animals (EU 2010/63).

### Cell cytometry and FACS

Testis samples (∼30 mg) employed for FACS were collected from seabream males showing >80% of motile and progressive spermatozoa, and more than 2 min of motility duration. Biopsies were cut into small pieces of ∼1 g and treated with 0.2% collagenase (Merck type 1A) for 1 h under agitation in non-activating medium (NAM; in mg/ml: 3.5 NaCl, 0.11 KCl, 1.23 MgCl_2_, 0.39 CaCl_2_, 1.68 NaHCO_3_, 0.08 glucose, 1 bovine serum albumine [BSA], pH 7.7; 280 mOsm) (*51*) supplemented with 200 μg/mL penicillin/streptomycin (Life Technologies Corp.). Samples were centrifuged at 200 × g for 1 min to remove cell aggregates, and the supernatant centrifuged again at 400 × g for 1 min to enrich in haploid cells. The cells were centrifuged at 400 × g for 5 min and the pellet resuspended in 1 ml NAM. The concentration of cells was determined by light microscopy and the ISASv1 software (Proiser), and this was adjusted to 150 x 10^6^ cells/ml. Cells were then stained with 200 nM of a solution of SYBR Green I (SGI) fluorescent nucleic acid stain (Molecular Probes, Life Technologies Corp.) for 45-60 min in the dark at room temperature, just prior to flow cytometry.

FACS was performed with a MoFlo XDP cell sorter (Beckman Coulter) equipped with three lasers (blue solid state of 488nm, red diode of 635nm, and argon ion UV laser of 351nm). Sterilized PBS served as the sheath fluid. The sorter was set in 4-way purify sort mode and with a flow sorting rate of ∼1500 events/s. The sorted population of HGC was collected in 4 ml of NAM in 15 ml tubes and centrifuged at 200 × g for 15 min. The resulting pellet was resuspended in 100 μl of NAM to obtain aliquots of 3 to 5 x 10^6^ cells, which were centrifuged again at 200 × g and frozen in liquid nitrogen and stored at -80°C.

### RNA extraction, library preparation, and sequencing

Total RNA from HGC (3 x 10^7^ cells) and SPZ_EJ_ (3-30 x 10^7^ cells) was extracted with the RNeasy Plus Mini Kit (Qiagen), and the purity and concentration of the extracted RNA was evaluated with the Agilent 2100 Bioanalyzer (Agilent Technologies). Four unstranded RNA libraries (replicates) for low-input RNA were constructed for each of the HGC and SPZ_EJ_ groups; each replicate being a pool of cells collected from three different males. The libraries from the total RNA were prepared following the SMARTseq2 protocol for low-input RNA (***Picelli et al., 2014***) with some modifications. Briefly, reverse transcription with 2 ng RNA was performed using SuperScript II (Invitrogen) in the presence of oligo-dT30VN (1µM; 5’-AAGCAGTGGTATCAACGCAGAGTACT_30_VN-3’), template-switching oligonucleotides (1 µM) and betaine (1 M). The cDNA was amplified using the KAPA Hifi Hotstart ReadyMix (Merck), 100 nM ISPCR primer (5’-AAGCAGTGGTATCAACGCAGAGT-3’) and 12 cycles of amplification. Following purification with Agencourt Ampure XP beads (1:1 ratio; Beckmann Coulter), product size distribution and quantity were assessed on a Bioanalyzer High Sensitvity DNA Kit (Agilent). The amplified cDNA (200 ng) was fragmented for 10 min at 55 °C using Nextera® XT (Illumina) and amplified for 12 cycles with indexed Nextera® PCR primers. The library was purified twice with Agencourt Ampure XP beads (0.8:1 ratio) and quantified on a Bioanalyzer using a High Sensitvity DNA Kit.

The libraries were sequenced on HiSeq2500 (Illumina) in paired-end mode with a read length of 2 x 76bp using TruSeq SBS Kit v4. We generated more than 30 million paired-end reads for each sample in a fraction of a sequencing v4 flow cell lane, following the manufacturer’s protocol. Image analysis, base calling and quality scoring of the run were processed using the manufacturer’s software Real Time Analysis (RTA 1.18.66.3) and followed by generation of FASTQ sequence files by CASAVA 1.8.

### Genome annotation

To improve the gilthead seabream reference genome (***Pauletto et al., 2018***) for the differential expression analysis, the genome was reannotated, and a *de novo* transcriptome assembly was generated from which those transcripts not present in the genome assembly were added to the analysis.

#### Genome reannotation

Repeats present in the seabream genome assembly were annotated with RepeatMasker v4-0-7 (http://www.repeatmasker.org) using the zebrafish repeat library included in RepeatMasker. The gene annotation was obtained by combining transcript alignments, protein alignments and *ab initio *gene predictions. First, the RNA-seq reads were aligned to the genome with STAR v-2.5.3a (***Dobin et al., 2013***). Subsequently, transcript models were generated using Stringtie v1.0.4 (***Pertea et al., 2015***) and PASA assemblies were produced with PASA v2.0.2 (***Haas et al., 2008***) by adding also the 114,155 *S. aurata* ESTs present in NCBI (October 2017). Secondly, the complete Actinopterygii proteomes were downloaded from Uniprot in October 2017 and aligned to the genome using Spaln v2.4.7 (***Iwata and Gotoh, 2012***). *Ab initio* gene predictions were performed on the repeat masked assembly with three different programs: GeneID v1.4 (***Parra et al., 2000***), Augustus v3.2.3 (***Stanke et al., 2006***) and Genemark-ES v2.3e (***Lomsadze et al., 2014***) with and without incorporating evidence from the RNA-seq data. The gene predictors were run with trained parameters for human except Genemark that runs on a self-trained manner. Finally, all the data was combined into consensus CDS models using EvidenceModeler-1.1.1 (***Haas et al., 2008***). Additionally, UTRs and alternative splicing forms were annotated through two rounds of PASA annotation updates. Functional annotation was performed on the annotated proteins with Blast2go (***Conesa et al., 2005***), using Blastp (***Altschul et al., 1990***) search against the nr database (March 2018) and Interproscan (***Jones et al., 2014***) to detect protein domains on the annotated proteins.

The annotation of ncRNAs was carried out by the following steps. First, the program cmsearch v1.1 (***Cui et al., 2016***) included in the Infernal software (***Nawrocki et al., 2015***) was run against the RFAM v12.0 database of RNA families (***Nawrocki et al., 2015***). The tRNAscan-SE v1.23 (***Chan and Lowe, 2019***) was also run in order to detect the transfer RNA genes present in the genome assembly. To detect the lncRNAs we selected those Pasa-assemblies that had not been included into the annotation of protein-coding genes in order to get all those expressed genes that were not translated into a protein. Finally, those PASA-assemblies without protein-coding gene annotation that were longer than 200 bp and whose length was not covered at least in an 80% by a small ncRNA were incorporated into the ncRNA annotation as lncRNAs. The resulting transcripts were clustered into genes using shared splice sites or significant sequence overlap as criteria for designation as the same gene.

#### Complementing the annotation with de novo assembled transcripts

The RNA-seq reads were assembled with Trinity v2.2.0 (***Grabherr et al., 2011***) allowing for trimming and normalization of the reads. Next, Rapclust v0.1 (***Trapnell et al., 2013***) was run, in which the process of pseudoalignment was first performed with Sailfish v0.10.0 (***Li et al., 2010***), and then Rapclust was used to cluster the assembled sequences into contained isoforms in order to reduce redundancy and to cluster together all the isoforms that are likely to belong to the same gene. For evaluation of the resulting transcriptomes we estimated their completeness with BUSCO v3.0.2 (Simao et al., 2015) using an Actinopterygii specific dataset of 4584 genes. After obtaining the reference transcriptome, open reading frames (ORFs) were annotated in the assembled transcripts with Transdecoder (***Haas et al., 2013***) and functional annotation was performed on the annotated proteins with Blast2GO, as described above. Finally, the assembled transcripts were mapped against the seabrem reference genome assembly with GMAP (Wu et al., 2005). Those transcripts for which less than 50% of their length aligned to the genome, and with a complete ORF and functional annotation, were added to the reference genome as separate annotated contigs.

### Differential expression analysis

RNA-seq reads were mapped against the improved version of the seabream reference genome with STAR v2.5.3a using ENCODE parameters for long RNAs. Genes were quantified with RSEM v1.3.0 (***Li and Dewey, 2011***) using the improved annotation. Sample similarities were inspected with a PCA using the top 500 most variable genes and the ’rlog’ transformation of the counts. Differential expression analysis was performed with DESeq2 v1.18 (***Love et al., 2014***) with default options, and genes with a false discovery rate (FDR) < 1% were considered significant. Heatmaps with the ‘rlog’ transformed counts of the DEGs were carried out with the ‘pheatmap’ R package. Venn diagrams and volcano plots were performed with the ‘VenDiagramm’ R package and ‘ggplot2’ R package, respectively.

### Gene classification, Ontology, and Pathway Analysis of DEGs

The GO enrichment of DEGs and signaling pathway analyses were performed using the PANTHER v14.1 Classification System and analysis tools (http://www.pantherdb.org/). GO terms and pathways with FDR < 0.05% were considered significant. Scattered plot of pathway analysis was carried out with ‘ggplot2’ R package. Functional categories classification were also done manually using the Uniprot database (https://www.uniprot.org/) and QuickGO browser (http://www.ebi.ac.uk/QuickGO). Interactome analyses were conducted using the STRING database v11.0b (*32*) with a high-confidence interaction score (0.9), and plots were performed using Cytoscape v3.8.2 (https://cytoscape.org/).

### In situ hybridization

Samples of seabream and zebrafish testis and ETDs were fixed in 4% paraformaldehyde (PFA) prepared in phosphate buffer saline (PBS: 137 mM NaCl, 2.7 mM KCl, 100 mM Na_2_HPO_4_, 2 mM KH_2_PO_4_, pH 7.4) overnight at 4°C. Samples were washed in PBS, dehydrated with increasing concentration of ethanol (50%, 70%, 95%, 100%) and xylene (100 %), and embedded in Paraplast^®^ (Merck). In situ hybridization was performed on 7-μm thick sections using digoxigenin-incorporated cRNA probes synthesized with SP6 and T7 RNA polymerases using the DIG RNA Labeling Mix (Merck 11277073910). Probes were specific for each target mRNA and did not share more than 35% identity between related transcripts (***Supplementary file 1***). Hybridization was performed at 45°C overnight with probe concentration at 2.5 µg/ml (*sbgnrh* and *sapdgfaa*, *-ba* and *-bb*) or 5 µg/ml (*sgnrh*, *sapdgfab*, *-c* and *-d*, and *drpdgfaa*, *-ab* and *-bb*). The post-hybridization washing included a first wash in 50% formamide in 2 x SSC at 45°C for 30 min, followed by two washes in 2 x SSC for 10 min at 45°C, and a final wash in 0.2 x SSC at 50°C. After blocking in TBST with 0.5% BSA, hybridized riboprobes were detected with alkaline phosphatase coupled rabbit anti-digoxigenin antibody (1:500; Merck 11093274910) for two hours at room temperature, and subsequent chromogenic revelation (NBT/BCIP Stock solution, Merck 11681451001). The reaction was stopped in distilled water and slides were mounted with Fluoromount™ aqueous mounting medium (Merck F4680).

### Immunofluorescence microscopy

Sorted germ cells and SPZ_EJ_ were processed as described previously (*51, 71*) and attached to UltraStick/UltraFrost Adhesion slides (Electron Microscopy Sciences). Samples were fixed in 4% PFA in PBS for 15 min before antigen retrieval in three consecutive 5-min incubations with boiling citrate (10 mM at pH 6), followed by triton X-100 (0.2% in PBS) for 15 min. After blocking for one hour in PBST with 5% normal goat serum (Merck G9023) and 0.1% BSA, antibodies were applied overnight at 4°C in a humidified chamber. The primary antibodies were α-tubulin (Merck T9026; 1:1,000), H3K9ac (Abcam ab4441; 1:1000), and Spo11 (Santa Cruz Biotechnology sc-33146; 1:1000). Anti-mouse or anti-rabbit IgG coupled with Alexa-555 (A-21422, Invitrogen, and AP510C, Merck, respectively) were applied for one hour at room temperature and cells were counterstained with 4′,6-diamidino-2-phenylindole dihydrochloride (DAPI; Merck G8294; 1:3000) before mounting with Fluoromount™.

The biopsies of testis and ETDs were fixed and processed as previously described (***Chauvigné et al., 2013***). Sections were permeabilized with 0.2% Triton X100 in PBS for 15 min, blocked with 5% normal goat serum, and subsequently incubated with affinity purified rabbit anti-GnRH (Merck, G8294, 1:400) in PBS+0.1% BSA overnight at 4°C. After washing, sections were incubated with a sheep anti-rabbit IgG antibody, Cy3 conjugate (Merck AP510C) for 2 h, the nuclei counterstained with DAPI (1:3000) for 3 min, and finally mounted with Fluoromount™.

### Sperm motility assays and in vitro incubation of SPZ_ED_

Freshly collected seabream SPZ_EJ_ and SPZ_ED_ were diluted 1:100 in NAM, whereas zebrafish SPZ_EJ_ and SPZ_ETD_ were no further diluted. Spermatozoa concentration and kinetic parameters were determined by computer-assisted sperm analysis (CASA) using the Integrated Semen Analysis System (ISASv1, Proiser) software as previously described (***Chauvigné et al., 2013***, ***2021***). The sperm kinetics analyses were run in triplicate (technical replicates) for each ejaculate. For seabream, the analyses were carried out on 4-8 different males (one ejaculate per male), whereas for zebrafish the analyses were done on three to four pools of three males each.

The SPZ_ED_ (10^7^ or 10^9^ cells/ml for zebrafish and seabream, respectively) were incubated in vitro in NAM (seabream) or modified SS300 medium for zebrafish (65 mM KCl, 62.5 mM NaCl, 2.35 mM CaCl_2_, 1 mM MgSO_4_, 6.5 mM MgCl_2_, 10 mM NaHCO_3_, 7 mM glucose, 30 mM Hepes-KOH pH 7.9, 0.015 mM BSA, pH 7.7; 330 Osm) in the presence of 100 nM of sbGnRH or sGnRH (Bachem, 4030832 and 4013835, respectively), 40 nM or rPDGF-BB (ThermoFisher Scientific PMG0044), or hormone vehicles (0.5% of water, or 0.8 mM acetic acid solution; controls). The incubations were carried out for 16-20 h at 16°C in a temperature-controlled incubator. After the incubation period, the sperm kinetic parameters were determined by CASA as above, and a subsample of SPZ_ED_ was frozen in liquid nitrogen and stored at -80°C until further RNA extraction. The effect of actinomycin D (Merck A9415) and chloramphenicol (Merck C1919) on motility was tested by preincubation of SPZ_ED_ with 100 µg/ml of the drugs for 1 h before addition of the hormones.

### Gene Expression Analyses

RT-PCR and qRT-PCR were carried out as previously described (***Chauvigné et al., 2013***, ***2014***), except that in this case the cDNA was synthesized from 1 µg (testis) or 13-20 ng (spermatozoa) of RNA using the AccuScript High-Fidelity 1st Strand cDNA Synthesis Kit (Agilent 200820) following the manufacturer’s instructions. For qRT-PCR, relative gene expression levels with respect to HGC or vehicle-treated SPZ_ED_ were determined by the 2^-ΔΔCt^ method, using glutathione-specific gamma-glutamylcyclotransferase 1 (*chac1*) or beta-actin (*bactin*) as reference genes. The analyses were done on three cDNAs synthesized from three different pools of three animals each, or on three to five cDNAs from different animals, using technical duplicates. Primer sequences are listed in ***Supplementary file 2***.

### Statistical analysis

Comparisons between two independent groups were made by the two-tailed unpaired Student’s *t*-test. The statistical significance among multiple groups was analyzed by one-way ANOVA, followed by the Tukey’s multiple comparison test, or by the non-parametric Kruskal-Wallis test and further Dunn’s test for nonparametric post hoc comparisons, as appropriate. Percentages were square root transformed previous analyses. Statistical analyses were carried out using the SigmaPlot software v12.0 (Systat Software Inc.) and GraphPad Prism v9.1.2 (226) (GraphPad Software). In all cases, statistical significance was defined as *P* < 0.05 (*), *P* < 0.01 (**), or *P* < 0.001 (***).

### Data availability

The RNA-seq datasets generated in this study have been submitted to Gene Expression Omnibus (GEO) database at the National Center for Biotechnology Information (NCBI) under accession no. GSE173088. Reannotation data are available at https://denovo.cnag.cat/Saurata. All other data generated or analysed during this study are included in the manuscript and supporting files.

## Acknowledgments

This work was supported by the Spanish Ministry of Science and Innovation (MICIN) Grant AGL2016-76802-R (to J.C.). F.C. and J.C.A. were supported, respectively, by the “Ramon y Cajal” programe (RYC-2015-17103) and a predoctoral (BES-2017-080778) contract from Spanish MICIN. A.E.C. was funded by ISCIII/ MICIN (PT17/0009/0019) and co-funded by FEDER, whereas R.N.F. was supported by the University of Bergen, Norway. We also acknowledge support of the Spanish MICIN through the Instituto de Salud Carlos III and to the EMBL partnership, the Centro de Excelencia Severo Ochoa, the Generalitat de Catalunya through the CERCA Programme, Departament de Salut and Departament d’Empresa i Coneixement, and funds from the European Regional Development Fund (Programa Operatiu FEDER de Catalunya 2014-2020) cofinanced by the Spanish MICIN (Programa Operativo FEDER Plurirregional de España (POPE) 2014-2020).

## Competing interests

The authors declare that no competing interests exist.

**Supplementary file 1.** Nucleotide sequences of the primers employed for ISH probe synthesis and alignment of probes.

**Supplementary file 2.** Nucleotide sequences of the primers employed for RT-PCR and qRT-PCR analyses.

---

**Figure 2-Supplement 1.**
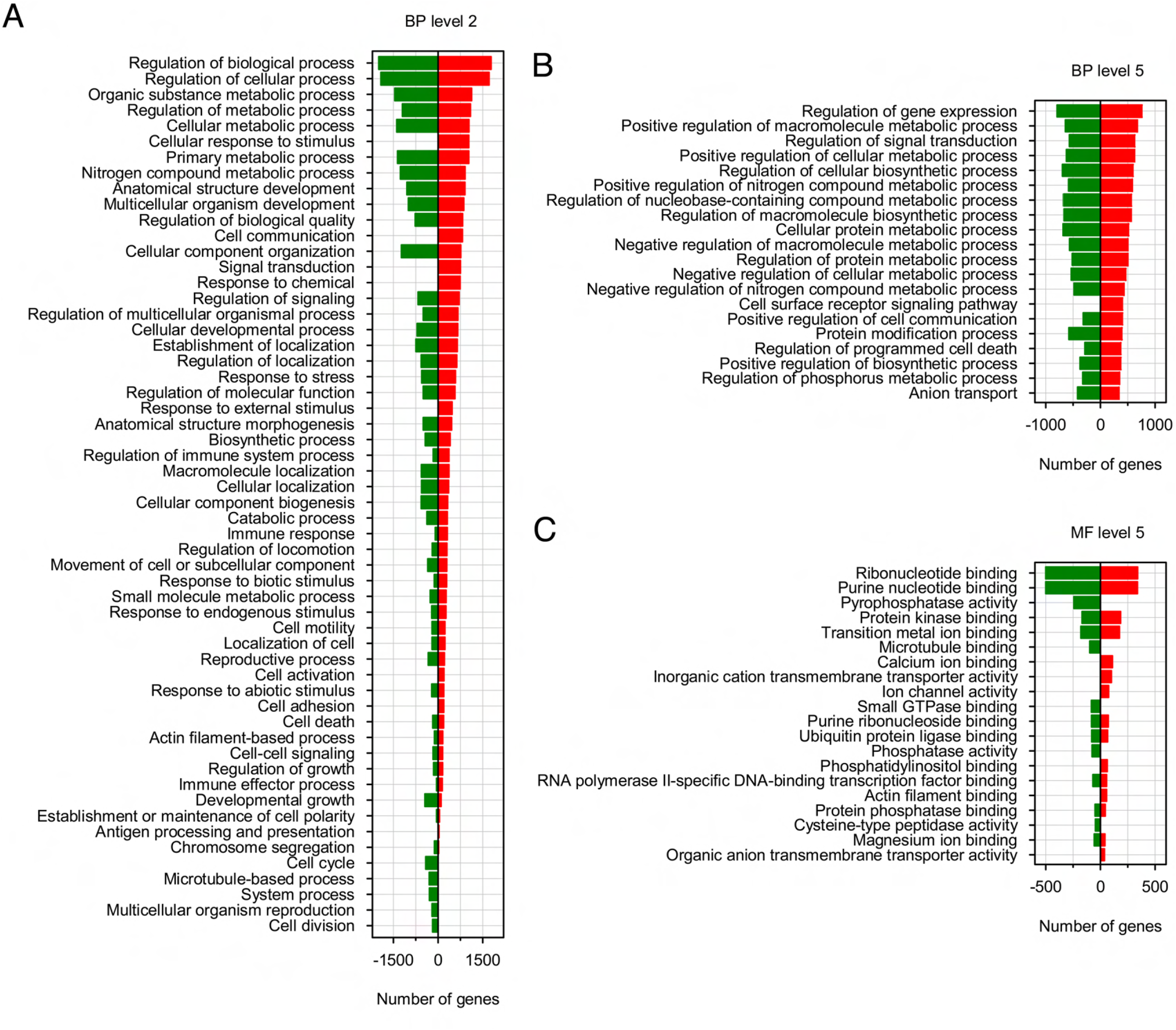
Gene ontology (GO) enrichment analysis of the DEGs during sperm differentiation and maturation. GO annotation of DEGs corresponding to biological process level 2 (A) and 5 (B), and molecular function level 5 (C). The horizontal axis displays the number of significant genes corresponding to each functional type, whereas the vertical axis displays the second level of GO annotation.

**Figure 3-Supplement 1.**
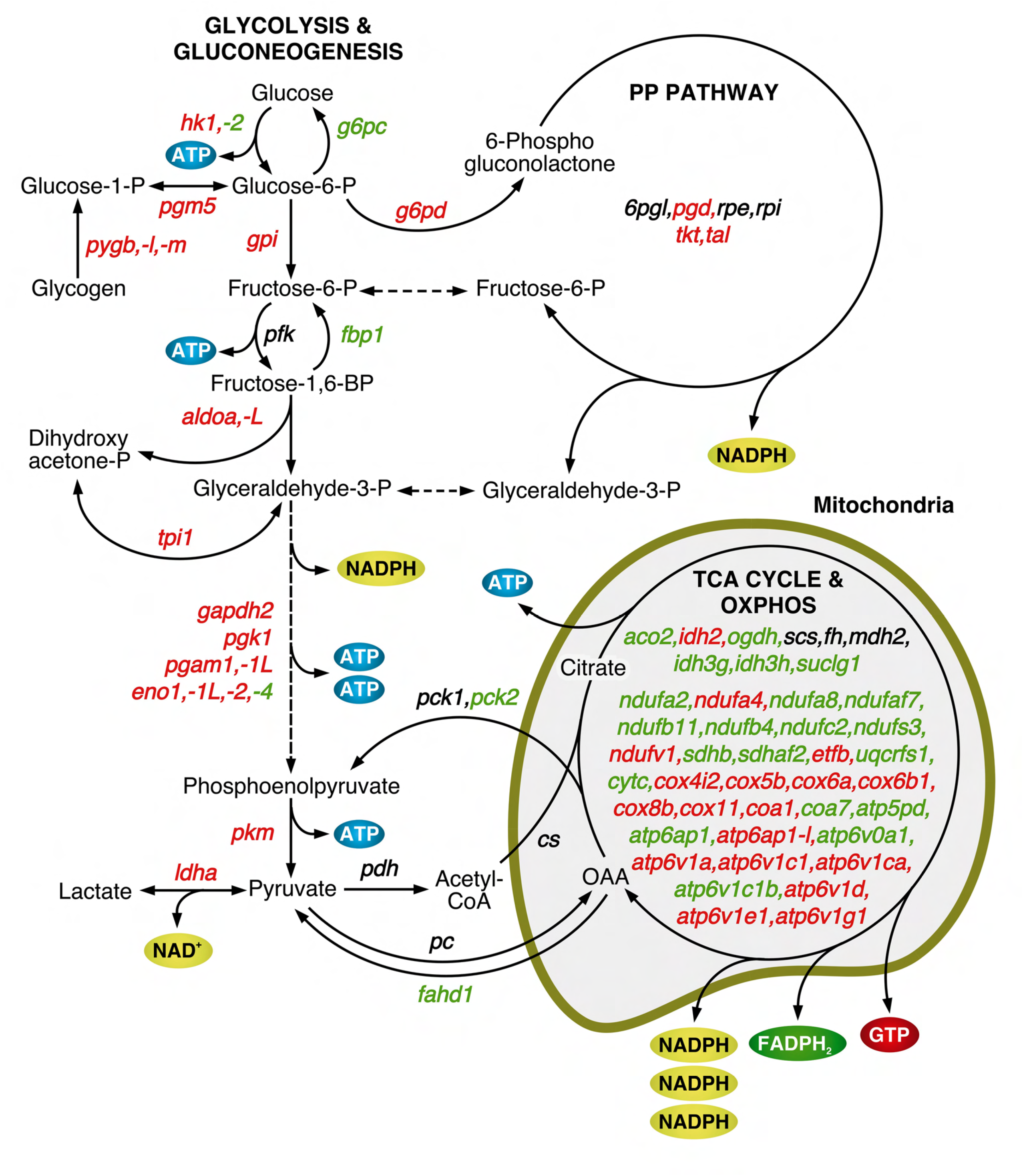
Mapping of DEGs coding for enzymes involved in respiratory pathways. Schematic diagram of the biochemical pathways of glycolysis/gluconeogenesis, penthose phosphate (PP) pathway, tricarboxylic acid (TCA) cycle and oxidative phophorylation (OXPHOS). Enzyme-coding DEGs in green and red color denotes downregulation and upregulation, respectively, whereas black color indicates no change in the expression levels.

**Figure 5-Supplement 1.**
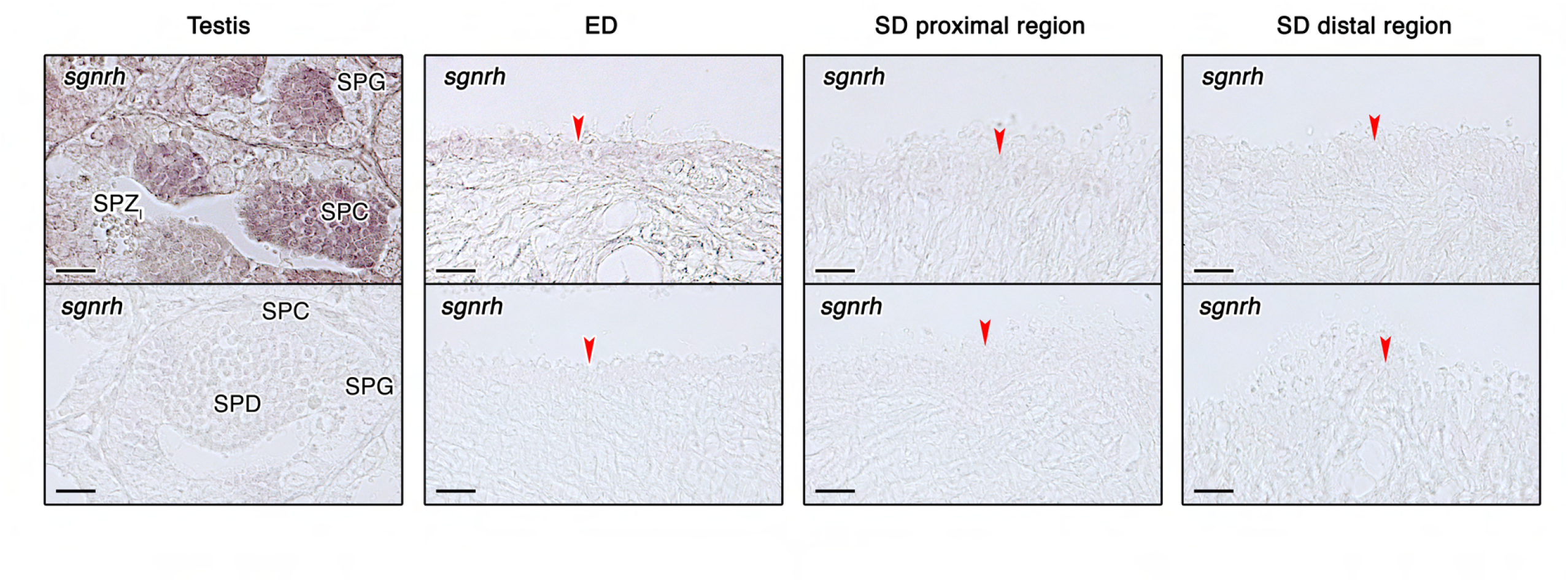
Localization of *sgnrh* transcripts in the sebream testis and efferent and spermatic ducts. Paraffin sections from the testis, efferent duct (ED) and two regions of the spermatic duc (SD) were hybridized with antisense DIG-labeled riboprobes specific for *sgnrh* or with specific sense probes (lower panels, negative controls). Scale bars, 50 µm. Abbreviations: SPG, spermatogonia; SPC, spermatocytes; SPD, spermatids; SPZ_I_, intratesticular spermatozoa; SC, Sertoli cells. The arrowheads indicate epithelial cells of the efferent and spermatic ducts.

**Figure 5-Supplement 2.**
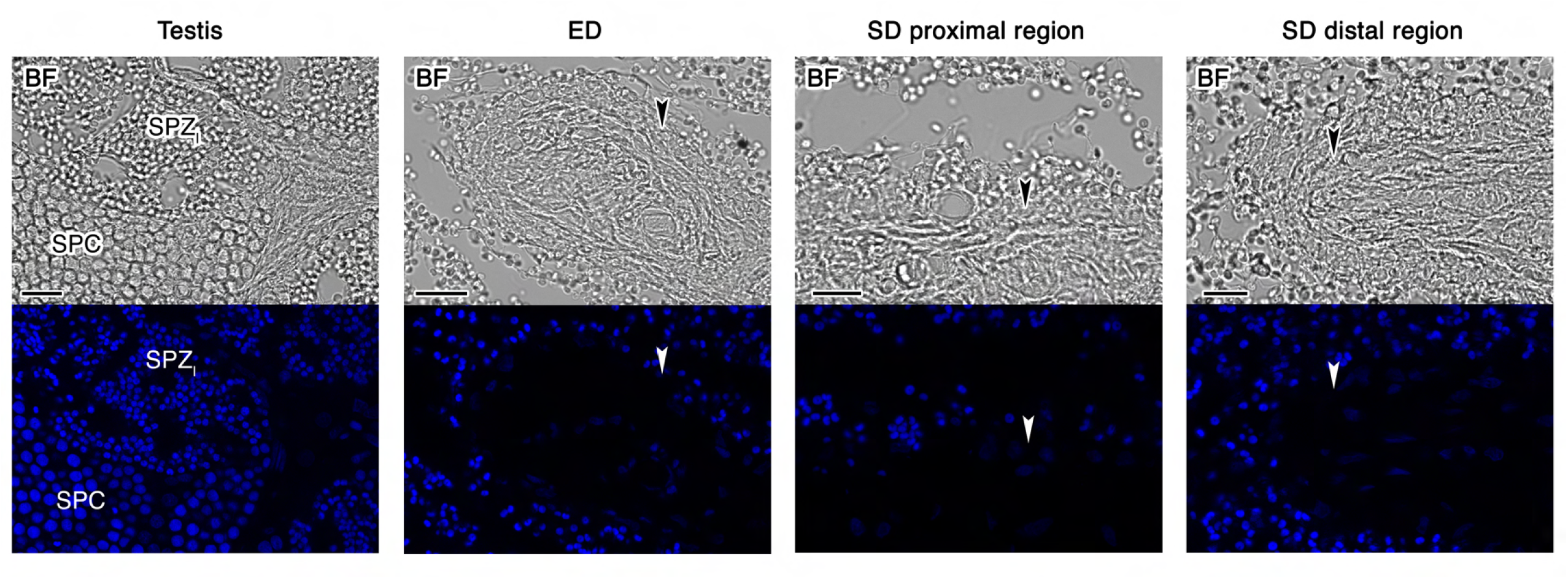
Control sections from the seabream testis, efferent duct (ED) and sperm duct (SD) incubated with the secondary antibody only. The upper panels show the brightfield (BF) images, whereas the lower panels show the epifluorescence images. Scale bars, 10 µm. Abbreviations: SPC, spermatocytes; SPZ_I_, intratesticular spermatozoa. The arrowheads indicate epithelial cells of the efferent and spermatic ducts.

**Figure 6-Supplement 1.**
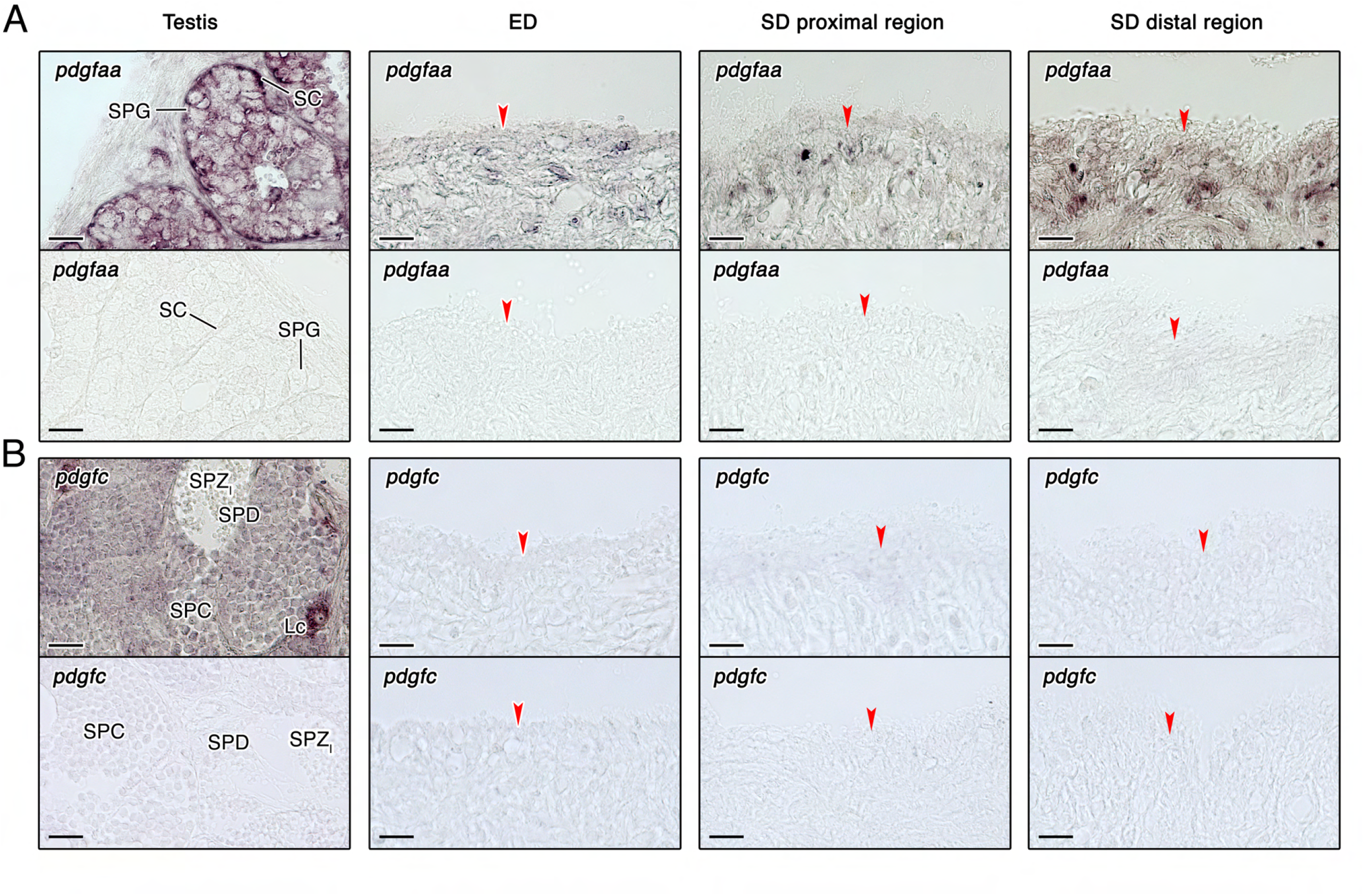
Localization of *pdgfaa* and *pdgfc* transcripts in the sebream testis and efferent and spermatic ducts. (A-B) Paraffin sections from the testis, efferent duct (ED) and two regions of the spermatic duc (SD) were hybridized with antisense DIG-labeled riboprobes specific for *pdgfaa* (A) and *pdgfc* (B) (upper panels) or with specific sense probes (lower panels, negative controls). Scale bars, 50 µm. Abbreviations: SPG, spermatogonia; SPC, spermatocytes; SPD, spermatids; SPZ_I_, intratesticular spermatozoa; SC, Sertoli cells. The arrowheads indicate epithelial cells of the efferent and spermatic ducts.

**Figure 7-Supplement 1.**
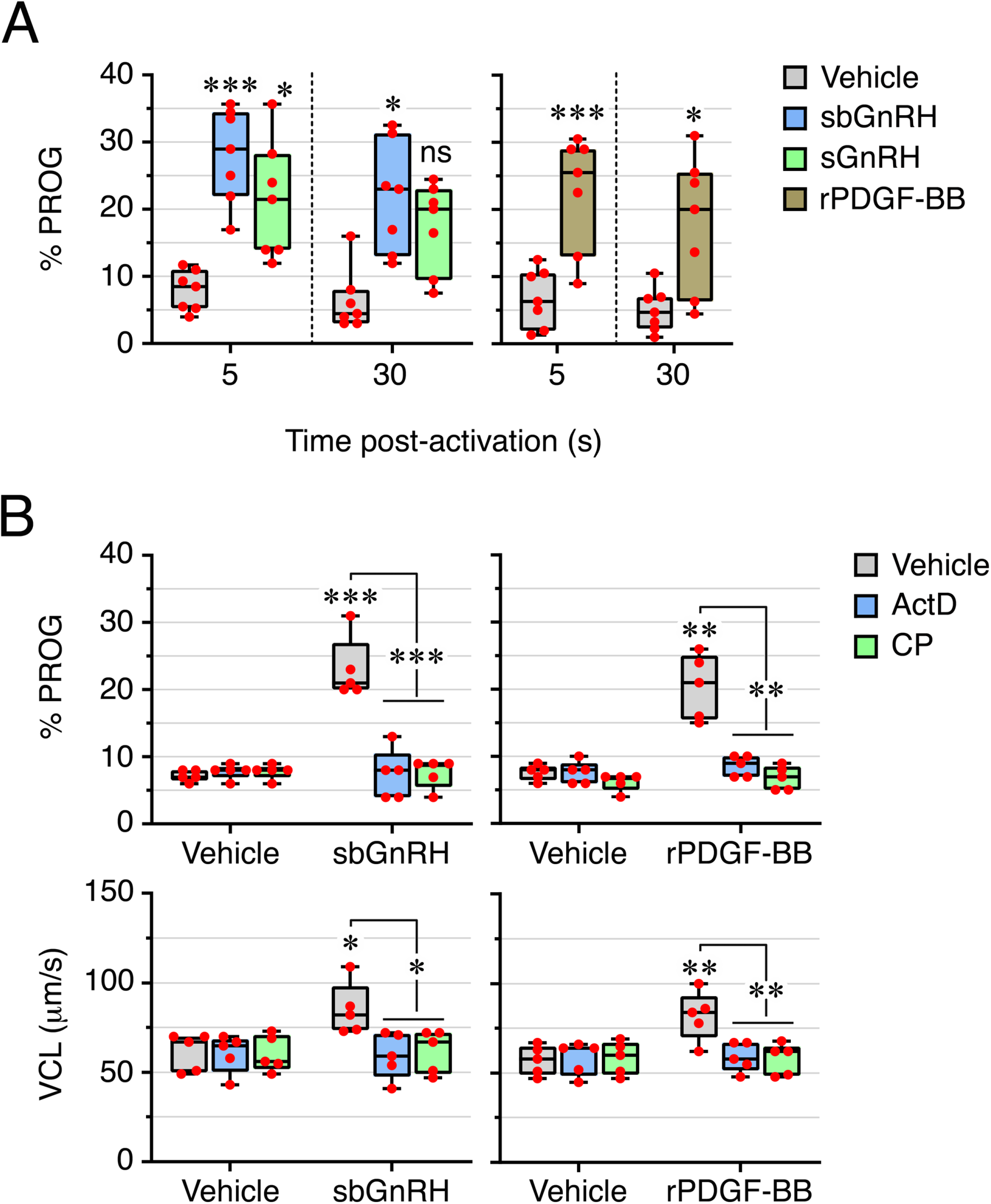
Sperm motion kinetics of seabream SPZ_ED_. (A) Percentage of progressivity (PROG) of SPZ_ED_ exposed to 100 nM of sbGnRH or sGnRH, 40 nM of recombinant PDGF (rPDGF-BB), or to each hormone vehicle, determined at 5 or 30 s postactivation. (B) Inhibition of PROG and curvilinear velocity (VCL) of SPZ_ED_ induced by sbGnRH and rPDGF-BB by 100 µg/ml actinomycin D (ActD) or chloramphenicol (CP) at 5 s postactivation. All data points are presented as box and whisker plots/scatter dots with horizontal line (inside box) indicating median and outliers. Data were statistically analyzed by one-way ANOVA. *, *P* < 0.05; **, *P* < 0.01; ***, *P* < 0.001; with respect to spermatozoa incubated with the hormone vehicles, or as indicated in brackets.

**Figure 8-Supplement 1.**
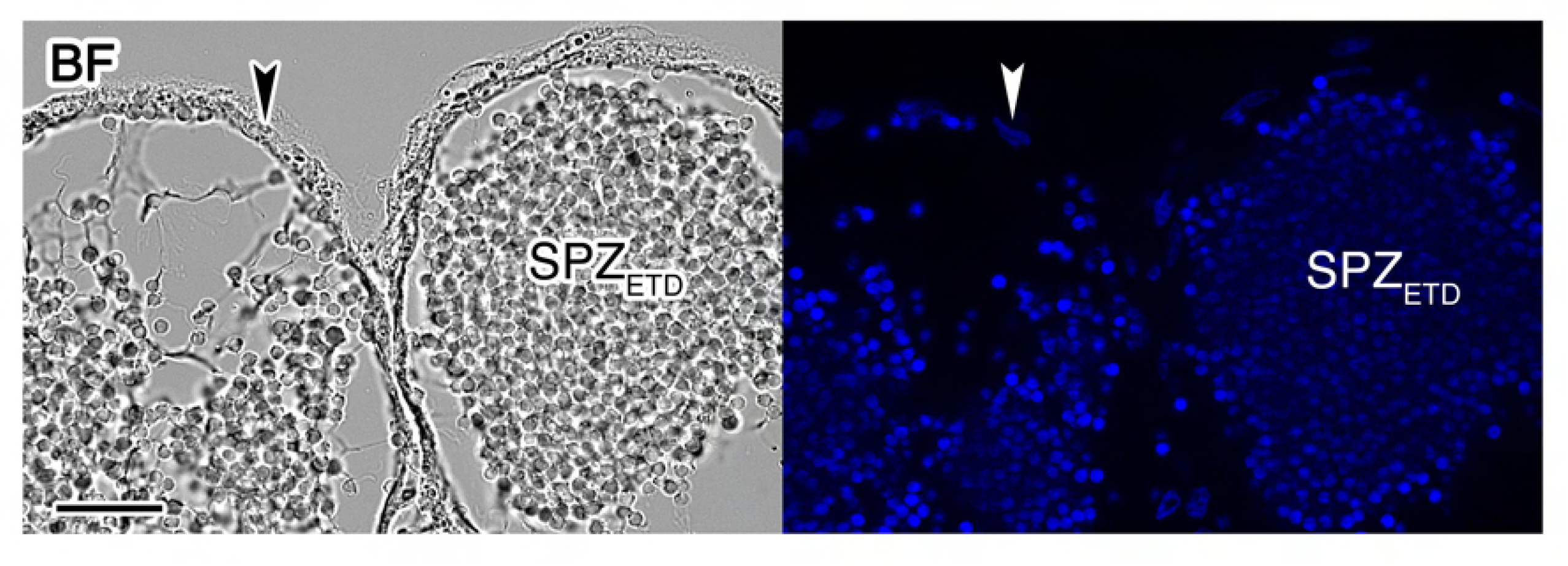
Control section of the zebrafish (ETDs) incubated with the secondary antibody only. The left panel shows the brightfield (BF) image, whereas the right panel show the epifluorescence image. The arrowheads indicate the ED epithelium. Scale bar, 200 µm. Abbreviations: SPZ_ETD_, sperm from the ETD.

**Figure 8-Supplement 2.**
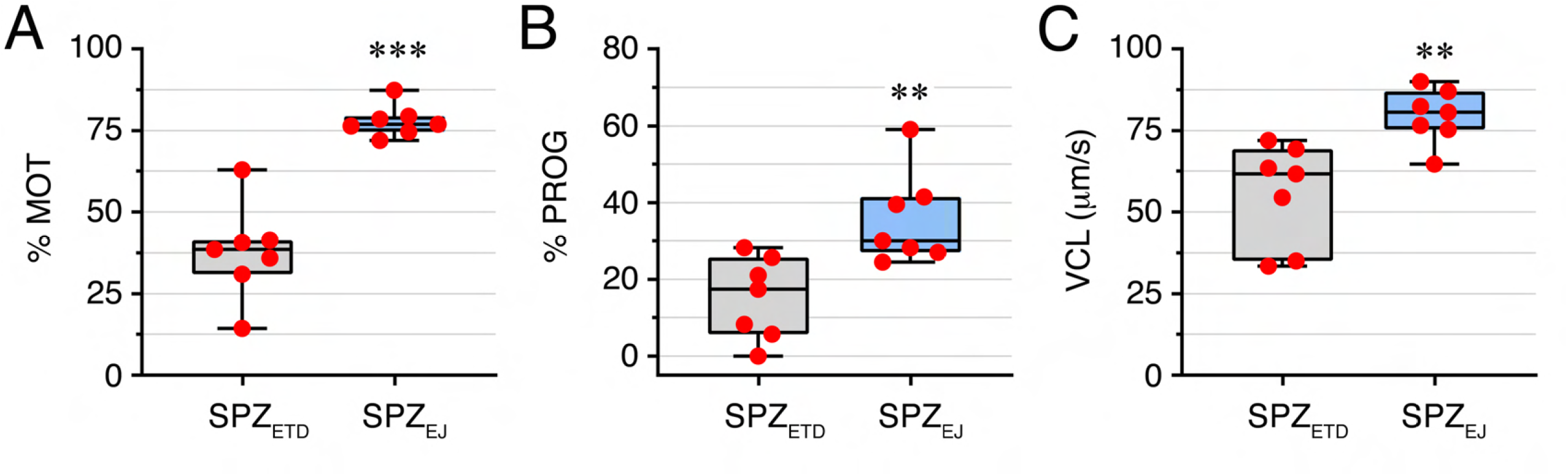
Kinematic properties of SPZ_ETD_ and SPZ_EJ_ from zebrafish. Percentage of motility (MOT) and progressivity (PROG), and curvilinear velocity (VCL), of zebrafish spermatozoa from extratesticular ducts (SPZ_ETD_) or ejaculated (SPZ_EJ_) determined at 5 s postactivation. All data points are presented as box and whisker plots/scatter dots with horizontal line (inside box) indicating median and outliers. One ejaculate from *n* = 7 males was measured. Data were statistically analyzed by an unpaired Student’s *t*-test. **, *P* < 0.01; ***, *P* < 0.001; with respect to SPZ_ED_.

